# *Tex19.1* Restricts LINE-1 Mobilisation in Mouse Embryonic Stem Cells

**DOI:** 10.1101/102442

**Authors:** Marie MacLennan, Marta García-Cañadas, Judith Reichmann, Elena Khazina, Carmen Salvador-Palomeque, Abigail R. Mann, Paula Peressini, Laura Sanchez, Christopher J. Playfoot, David Read, Chao-Chun Hung, Ragnhild Eskeland, Richard R. Meehan, Oliver Weichenrieder, Jose Luis GarcíaPérez, Ian R. Adams

**Author notes:** Correspondence to: JLGP or IRA. These authors contributed equally to this work (Marie MacLennan, Marta García-Cañadas). Current addresses: JR: EMBL Heidelberg, Meyerhofstraße 1, 69117 Heidelberg, Germany. CSP: Mater Research Institute, University of Queensland, TRI Building, Woolloongabba QLD 4102, Australia.

## Abstract

Mobilisation of retrotransposons to new genomic locations is a significant driver of mammalian genome evolution. In humans, retrotransposon mobilisation is mediated primarily by proteins encoded by LINE-1 (L1) retrotransposons, which mobilise in pluripotent cells early in development. Here we show that TEX19.1, which is induced by developmentally programmed DNA hypomethylation, can directly interact with the L1-encoded protein L1-ORF1p, stimulate its polyubiquitylation and degradation, and restrict L1 mobilisation. We also show that TEX19.1 likely acts, at least in part, through promoting the activity of the E3 ubiquitin ligase UBR2 towards L1-ORF1p. Moreover, we show that loss of *Tex19.1* increases L1-ORF1p levels and mobilisation of L1 reporters in pluripotent mouse embryonic stem cells implying that *Tex19.1* prevents new retrotransposition-mediated mutations from arising in the germline genome. These data show that post-translational regulation of L1 retrotransposons plays a key role in maintaining trans-generational genome stability in the epigenetically dynamic developing mammalian germline.

## Introduction

Retrotransposons are mobile genetic elements that comprise around 40% of mammalian genomes (Beck et al. 2011; Hancks and Kazazian 2016; Richardson et al. 2014a). Retrotransposons are a source of genetic variation that shape genome evolution and mammalian development, but their mobilisation can also cause mutations associated with a variety of genetic diseases and cancers (Beck et al. 2011; Hancks and Kazazian 2016; Richardson et al. 2014a; Garcia-Perez et al. 2016). New retrotransposition events are estimated to occur in around 1 in every 20 human births, and represent around 1% of genetic disease-causing mutations in humans (Kazazian 1999; Hancks and Kazazian 2016). Retrotransposons are classified into three major types depending on their genomic structure: LINEs (long interspersed nuclear elements), SINEs (short interspersed nuclear elements) and LTR (long terminal repeat) retrotransposons. In humans, all new retrotransposition events are catalysed by LINE-1 (L1) elements. Active L1s encode two proteins required for retrotransposition: ORF1p is an RNA binding protein with nucleic acid chaperone activity, and ORF2p is a multidomain protein with reverse transcriptase and endonuclease activities (Beck et al. 2011; Hancks and Kazazian 2016; Richardson et al. 2014a). Both these proteins interact directly or indirectly with various cellular factors and are incorporated into ribonucleoprotein particles (RNPs) along with L1 RNA (Beck et al. 2011; Goodier et al. 2013; Hancks and Kazazian 2016; Richardson et al. 2014a; Taylor et al. 2013). While these proteins exhibit a *cis*-preference to bind to and catalyse mobilisation of their encoding mRNA, they can act in *trans* on other RNAs, including those encoded by SINEs (Kulpa and Moran 2006; Wei et al. 2001; Dewannieux et al. 2003). Human L1 also encodes a *trans*-acting protein, ORF0, that stimulates retrotransposition, although its mechanism of action is currently poorly understood (Denli et al. 2015). Regulating the activity of these L1-encoded proteins will impact on the stability of mammalian genomes and the incidence of genetic disease.

Regulating retrotransposon activity is particularly important in the germline as *de novo* retrotransposon integrations that arise in these cells can be transmitted to the next generation (Crichton et al. 2014). The mammalian germline encompasses lineage-restricted germ cells including primordial germ cells, oocytes, and sperm, and their pluripotent precursors in early embryos (Ollinger et al. 2010). L1 mobilisation may be more prevalent in pluripotent cells in pre-implantation embryos rather than in lineage-restricted germ cells (Kano et al. 2009), and regulation of L1 activity in the pluripotent phase of the germline cycle is therefore likely to have a significant effect on trans-generational genome stability. Repressive histone modifications and DNA methylation typically suppress transcription of retrotransposons in somatic mammalian cells (Beck et al. 2011; Hancks and Kazazian 2016; Richardson et al. 2014a; Crichton et al. 2014), but many of these transcriptionally repressive marks are globally removed during pre-implantation development and during fetal germ cell development in mice (Hajkova et al. 2008; Popp et al. 2010; Santos et al. 2002; Fadloun et al. 2013). DNA methylation in particular plays a key role in transcriptionally repressing L1 in the germline (Bourc’his and Bestor 2004), and it is not clear how L1 activity is controlled in pluripotent cells and fetal germ cells while they are DNA hypomethylated.

In fetal germ cells, loss of DNA methylation correlates with relaxed transcriptional suppression of retrotransposons (Molaro et al. 2014), but also induces expression of methylation-sensitive germline genome-defence genes that have roles in post-transcriptionally repressing these elements (Hackett et al. 2012). The methylation-sensitive germline genome-defence genes include components of the PIWI-piRNA pathway. This pathway promotes *de novo* DNA methylation of retrotransposons in male germ cells, cleaves retrotransposon RNAs, and may also suppress retrotransposon translation (Fu and Wang 2014; Xu et al. 2008). However, while mice carrying mutations in the PIWI-piRNA pathway can strongly de-repress L1-encoded RNA and protein during spermatogenesis (Aravin et al. 2007; Carmell et al. 2007), increased L1 mobilisation has not yet been reported in these mutants.

Indeed, the level of L1 expression at different stages of the germline cycle does not completely correlate with the ability of L1 to mobilise and post-translational control mechanisms have been proposed to restrict the ability of L1 to mobilise in the mouse germline (Kano et al. 2009). However, the molecular identities of these post-translational L1 restriction mechanisms have not yet been elucidated.

Here we show that one of the genes induced in response to programmed DNA hypomethylation in the mouse germline, *Tex19.1*, regulates L1-ORF1p levels and mobilisation of L1 reporters. We show that mouse TEX19.1, and its human ortholog, physically interact with L1-ORF1p, and regulate L1-ORF1p abundance through stimulating its polyubiquitylation and proteasome-dependent degradation. We show that TEX19.1 likely controls L1-ORF1p abundance in concert with UBR2, an E3 ubiquitin ligase that we show also physically interacts with and regulates L1-ORF1p levels *in vivo*. As anticipated from our analysis, we show that loss of *Tex19.1* results in increased L1-ORF1p abundance and increased mobilisation of L1 reporters in pluripotent mouse embryonic stem cells, suggesting that *Tex19.1* functions as a post-translational control mechanism to restrict L1 mobilisation in the developing germline.

## Results

### L1-ORF1p Abundance Is Post-Transcriptionally Regulated By *Tex19.1* In Mouse Germ Cells

We have previously shown that programmed DNA hypomethylation in the developing mouse germline induces expression of a group of genes that are involved in suppressing retrotransposon activity (Hackett et al. 2012). One of the retrotransposon defence genes induced in response to programmed DNA hypomethylation, *Tex19.1*, suppresses specific retrotransposon transcripts in spermatocytes (Öllinger et al. 2008; Reichmann et al. 2012), however its direct mechanism of action remains unclear. *Tex19.1* is expressed in germ cells, pluripotent cells and the placenta and is one of two *TEX19* orthologs generated by a rodent-specific gene duplication (Kuntz et al. 2008; Öllinger et al. 2008). These mammal-specific proteins have no functionally characterised protein motifs or reported biochemical activity, but mouse TEX19.1 is predominantly cytoplasmic in the germline (Öllinger et al. 2008; Yang et al. 2010). We therefore investigated if *Tex19.1* has post-transcriptional effects on cytoplasmic stages of the retrotransposon life cycle. Since *Tex19.1*^*−/−*^spermatocytes have defects in meiosis that induce spermatocyte death (Öllinger et al. 2008), we analysed mouse L1 ORF1p (mL1-ORF1p) expression in prepubertal testes during the first wave of spermatogenesis before any increased spermatocyte death is evident (Öllinger et al. 2008). Western blotting showed that P16 *Tex19.1*^*−/−*^ testes have elevated levels of mL1-ORF1p (Figure 1A), even though L1 RNA levels do not change (Figure 1B) (Öllinger et al. 2008; Reichmann et al. 2012) suggesting that *Tex19.1* negatively regulates mL1-ORF1p post-transcriptionally in male germ cells. Immunostaining of P16 testes showed that while mL1-ORF1p is expressed in meiotic spermatocytes in control mice, consistent with previous reports (Soper et al. 2008; Branciforte and Martin 1994), that mL1-ORF1p immunostaining is elevated in this cell type in *Tex19.1*^*−/−*^ mice (Figure 1C). Thus, distinct from its role in transcriptional regulation of retrotransposons (Öllinger etal. 2008; Reichmann et al. 2012; Crichton et al. 2017a; Reichmann et al. 2013), *Tex19.1* appears to have a role in post-transcriptionally suppressing mL1-ORF1p abundance in meiotic spermatocytes.

**Figure 1.**
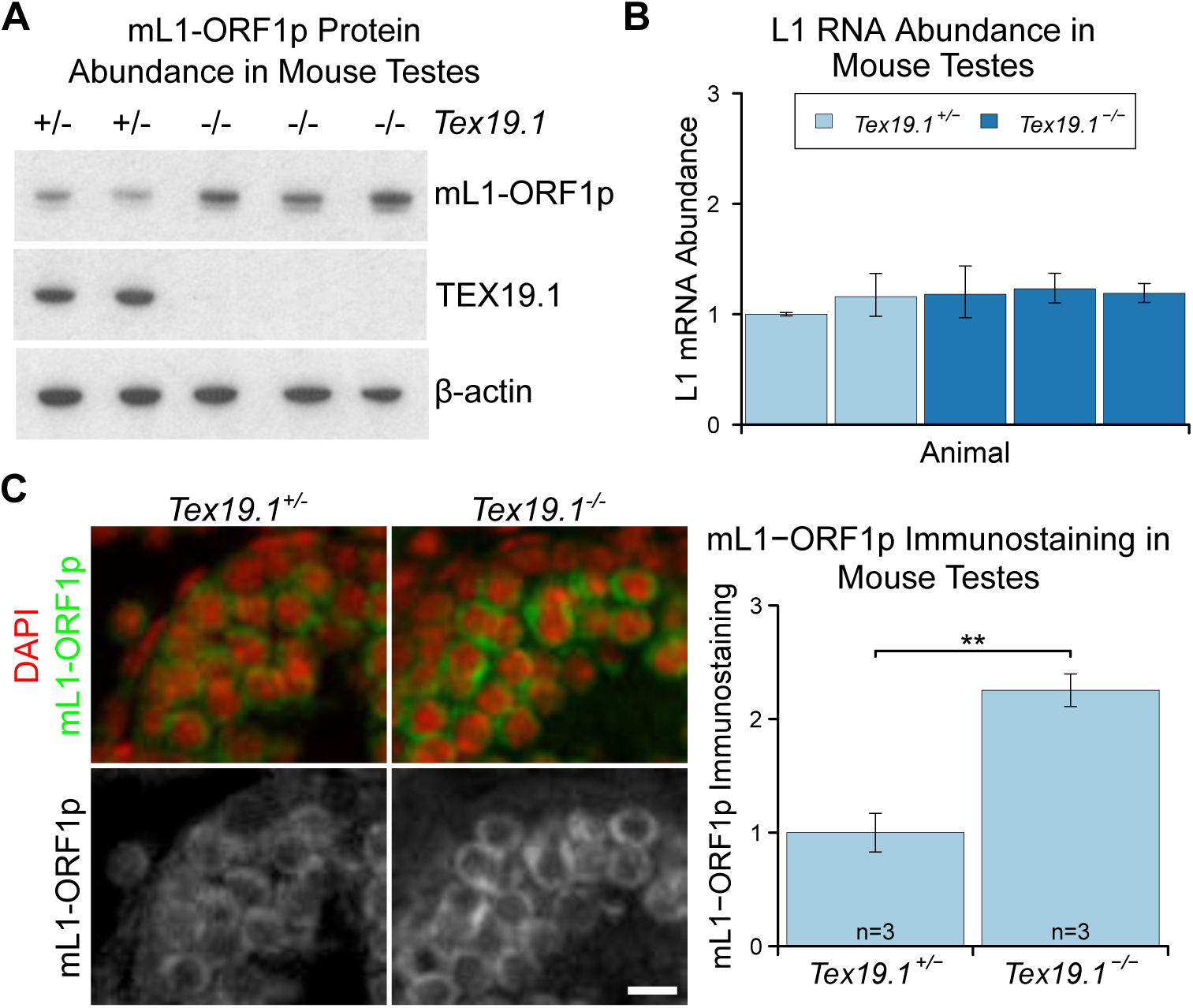
mL1-ORF1p Is Post-Transcriptionally Regulated By *Tex19.1* In Mouse Germ Cells. A. Western blot for mL1-ORF1p in *Tex19.1*^*+/−*^ and *Tex19.1*^*−/−*^littermate P16 mouse testes. β-ctin is a loading control. B. qRT-PCR for L1 RNA in testes from the same animals analyzed in panel A. Expression relative to β-actin was normalised to a *Tex19.1*^*+/−*^ control animal. Error bars indicate SEM for three technical replicates. C. Immunostaining for mL1-ORF1p (green) in *Tex19.1*^*+/−*^ and *Tex19.1*^*−/−*^P16 mouse testis sections. Nuclei are counterstained with DAPI (shown as red). Scale bar,10 μm. Anti-mL1-ORF1p immunostaining per unit area was quantified for three animals for each genotype, and normalised to the mean for *Tex19.1*^*+/−*^ animals. Means ± SEM (1±0.17 and 2.25±0.14 for *Tex19.1*^*+/−*^ and *Tex19.1*^*−/−*^ respectively) are indicated; ** p<0.01 (*t*-test).

### TEX19.1 Interacts With Multiple Components Of The Ubiquitin-Proteasome System

Post-transcriptional control of protein abundance can occur through regulation of mRNA translation or protein stability. To investigate whether TEX19.1 might be involved in one of these processes we attempted to identify RNAs or proteins that interact with TEX19.1. In contrast to the PIWI proteins MILI and MIWI (Grivna et al. 2006; Unhavaithaya et al. 2009), oligo(dT) pull-downs from mouse testicular lysate suggest that TEX19.1 is not physically associated with RNA in this tissue (Supplementary Figure S1A) and neither is TEX19.1 enriched in testicular polysome fractions containing actively translating mRNAs (Supplementary Figure S1B). In addition, the increase in mL1-ORF1p abundance in *Tex19.1*^*−/−*^ testes is not accompanied by an increase in L1 RNA abundance in polysomes (Supplementary Figure S1C). Therefore the increase in mL1-ORF1p abundance in *Tex19.1* ^*−/−*^ testes does not appear to reflect a direct role for TEX19.1 in regulation of L1 translation. However, mass spectrometry of TEX19.1-YFP immunoprecipitates (IPs) from stably expressing mouse embryonic stem cells (ESCs) revealed co-IP of multiple components of the ubiquitin-proteasome system (Figure 2A, Figure 2B, Supplementary Table 1). TEX19.1-YFP IPs contained a strong co-immunoprecipitating band of approximately stoichiometric abundance to TEX19.1-YFP which was identified as UBR2, a RING domain E3 ubiquitin ligase and known interacting partner for TEX19.1 (Yang et al. 2010) (Figure 2A, Figure 2B, Supplementary Figure S2A). The identification of the only known interacting partner for TEX19.1 in this co-IP suggests that the TEX19.1-YFP construct used in this experiment recapitulates interactions relevant for endogenous TEX19.1. Indeed, all detectable endogenous TEX19.1 in ESCs co-fractionates with UBR2 in size exclusion chromatography (Figure 2C), consistent with TEX19.1 existing in a stable heteromeric complex with UBR2 *in vivo*. *Ubr2* has previously been shown to be required for TEX19.1 protein stability in mouse testes (Yang et al. 2010) which, in combination with the co-fractionation and stoichiometric abundance of these proteins in the ESC IPs, suggests that any TEX19.1 protein not associated with UBR2 may be unstable and degraded. TEX19.1-YFP also co-IPs with additional components of the ubiquitin-proteasome system including UBE2A/B, an E2 ubiquitin-conjugating enzyme and cognate partner of UBR2 (Kwon et al. 2003), and a HECT-domain E3 ubiquitin ligase, HUWE1 (Figure 2B, Supplementary Table 1). The physical associations between TEX19.1 and multiple components of the ubiquitin-proteasome system suggest that the post-transcriptional increase in mL1-ORF1p abundance in *Tex19.1* ^*−/−*^ testes might reflect a role for TEX19.1 in regulating degradation of mL1-ORF1p.

**Figure 2.**
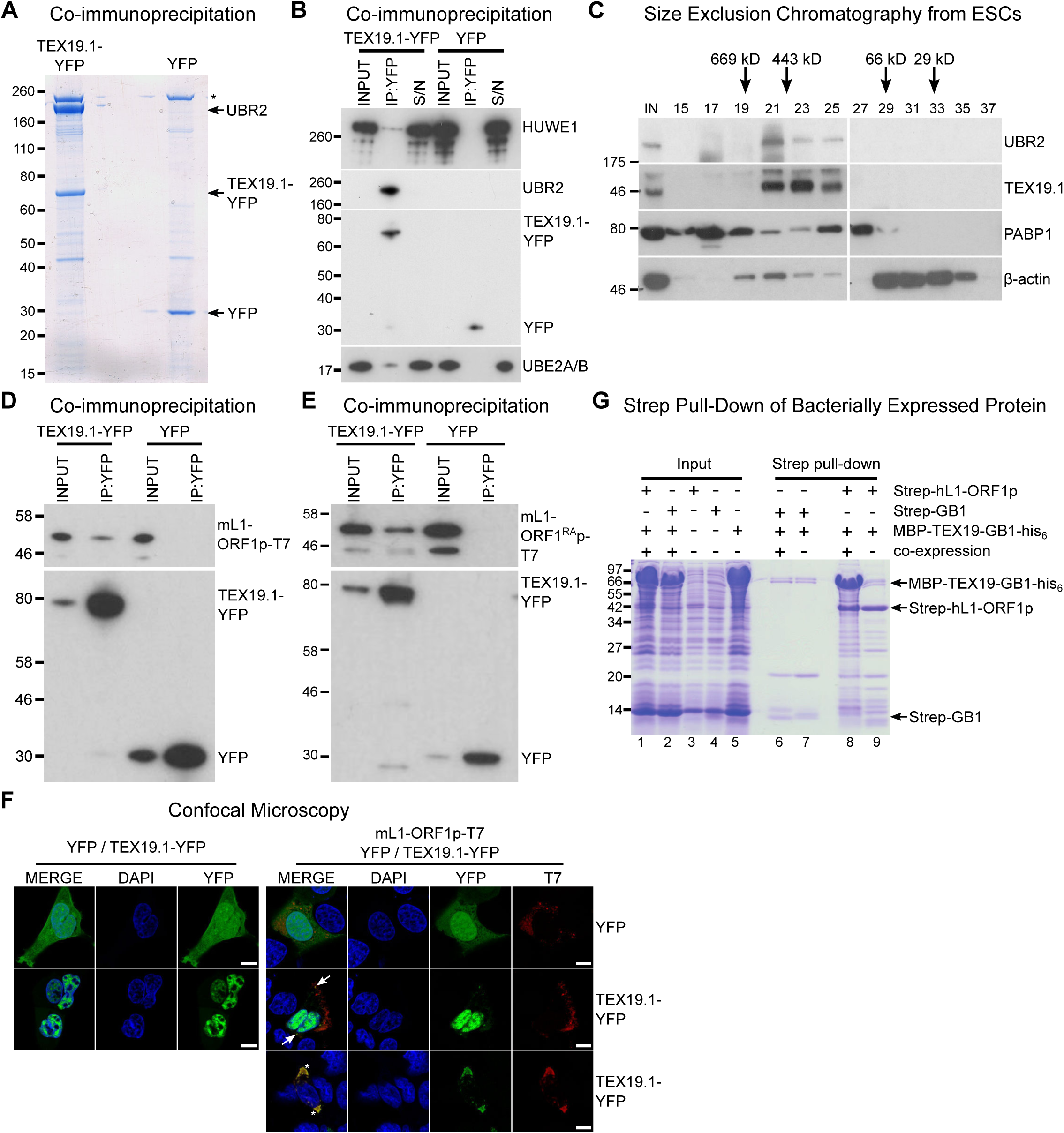
*TEX19* Orthologs Physically Interact With The Components Of The Ubiquitin Proteasome System And With L1-ORF1p. **A.** Colloidal blue-stained cytoplasmic anti-YFPimmunoprecipitates from mouse ESCs stably expressing TEX19.1-YFP or YFP. Mass spectrometry identities of major bands are indicated, and a non-specific band marked with an asterisk. **B.** Western blots for ubiquitin-proteasome system components in anti-YFP immunoprecipitates (IPs) from panel A. Anti-YFP IP inputs, IPs and IP supernatants (S/N) were blotted with indicated antibodies. **C.** Size exclusion chromatography of cytoplasmic extract from ESCs showing elution of endogenous TEX19.1 and UBR2. PABP1 and β-actin are included as controls. Input (IN) sample is also shown, and eluted fraction numbers and the positions of molecular weight markers in kD are indicated. **D, E.** IPs from HEK293T cells co-transfected with mL1-ORF1p-T7 constructs and either TEX19.1-YFP or YFP and Western blotted with indicated antibodies. The mutant mL1-ORF1^RA^p in panel E has a reduced binding affinity for RNA (Kulpa and Moran 2005; Martin et al. 2005). **F.** Subcellular localisation of TEX19.1-YFP in the presence and absence of mL1-ORF1p-T7. U2OS cells were transiently transfected with TEX19.1-YFP or YFP expression constructs with or without a plasmid expressing mL1-ORF1p-T7 (pCEPL1SM-T7), then stained with anti-T7 antibodies, and with DAPI to detect DNA. Around 40% of cells exhibit partial co-localization of a subset of small cytoplasmic foci of mL1-ORF1p-T7 with TEX19.1-YFP (arrows). In more extreme examples, large cytoplasmic aggregates of mL1-ORF1p-T7 extensively co-localise with TEX19.1-YFP (asterisks). Scale bars 10 μm. **G.** Strep pull-down assays from *E. col*i lysates. Double-tagged human TEX19 was either co-expressed with Strep-tagged hL1-ORF1p (lane 8) or added after hL1-ORF1p immobilisation on Strep-Tactin beads (lane 9). Strep-GB1 served as a control (lanes 6 and 7).

### TEX19.1 Orthologs Directly Interact With L1-ORF1p

We next tested if TEX19.1 might also interact with mL1-ORF1p. Although we did not identify any mL1-ORF1p peptides in the mass spectrometry analysis of TEX19.1-YFP IPs from ESCs, we did identify a single hL1-ORF1p peptide in similar IPs from stable TEX19.1-YFP expressing HEK293T cells. Interactions between E3 ubiquitin ligases and their substrates are expected to be transient and weakly represented in IP experiments, therefore we tested directly whether TEX19.1-YFP and epitope-tagged mL1-ORF1p interact by co-expressing these proteins in HEK293T cells and immunoprecipitating either TEX19.1-YFP or epitope-tagged mL1-ORF1p. Both IPs revealed weak reciprocal interactions between TEX19.1-YFP and epitope-tagged mL1-ORF1p (Figure 2D, Supplementary Figure S2C). Although human TEX19 is significantly truncated relative to its mouse ortholog, the physical interaction between TEX19 and L1-ORF1p is conserved in humans (Supplementary Figure S2D, Supplementary Figure S2E).

We next tested whether the biochemical interaction between TEX19.1-YFP and mL1-ORF1p-T7 is reflected by co-localisation of these proteins. TEX19.1 is predominantly cytoplasmic in ES cells and in germ cells (Öllinger et al. 2008; Yang et al. 2010), but in the hypomethylated placenta and when expressed in somatic cell lines TEX19.1 can localise to the nucleus (Kuntz et al. 2008; Reichmann et al. 2013). The context-dependent localisation of TEX19.1 suggests that TEX19.1-interacting proteins in ES cells and germ cells could retain this protein in the cytoplasm in these cell types. L1-ORF1p has been reported to form cytoplasmic aggregates that co-localise with stress granule markers (Doucet et al. 2010), therefore we tested whether co-expression of L1-ORF1p and TEX19.1 might localise TEX19.1 to these L1-ORF1p-containing aggregates. As expected, confocal microscopy showed that TEX19.1-YFP localises to the nucleus when expressed in U2OS cells, however co-expression with mL1-ORF1p-T7 resulted in partial co-localisation of both these proteins in cytoplasmic aggregates in around 40% of cells (Figure 2F). In more extreme examples, co-expression of mL1-ORF1p-T7 re-localised all detectable TEX19.1-YFP out of the nucleus and into cytoplasmic aggregates (Figure 2F). These co-localisation data are consistent with the co-IP data suggesting that TEX19.1-YFP and mL1-ORF1p-T7 physically interact.

A number of host factors have been shown to associate with L1-ORF1p, although many of these interactions are indirect and mediated by RNA, likely reflecting interactions within the L1 RNP (Goodier et al. 2013; Taylor et al. 2013). However, TEX19.1-YFP also interacts with a mutant allele of mL1-ORF1p which has severely impaired binding to RNA and impaired mobilisation (Kulpa and Moran 2005; Martin et al. 2005) (Figure 2E, Supplementary Figure S2F), suggesting that the interaction between TEX19.1-YFP and mL1-ORF1p is RNA-independent and could potentially be direct. We therefore tested whether bacterially expressed TEX19 might interact with bacterially expressed hL1-ORF1p. Co-expression of double-tagged human MBP-TEX19-GB1-His_6_ with Strep-tagged human L1-ORF1p (Strep-hL1-ORF1p) in bacteria resulted in a strong interaction betweenthese proteins, and isolation of a stable TEX19-hL1-ORF1p complex (Figure 2G). Taken together, the co-IPs, co-localisation and isolation of a TEX19-L1-ORF1p from bacterially expressed proteins suggest that TEX19 directly interacts with L1-ORF1p and, to our knowledge, represents the first example of a mammalian host protein that directly binds to a protein encoded by L1 retrotransposons.

### *Tex19.1* Orthologs Stimulate Polyubiquitylation and Degradation of L1-ORF1p

The strong interaction between TEX19 and hL1-ORF1p seen with bacterially-expressed proteins contrasts with weaker interactions detected in HEK293T cells. However, it is possible that the difference in the strength of these interactions reflects the presence of UBR2 in HEK293T cells, which allows a TEX19-UBR2 complex to assemble and transiently interact with hL1-ORF1p to catalyse its ubiquitylation and subsequent degradation. We therefore investigated if L1-ORF1p is ubiquitylated and degraded by the proteasome, and whether this might be stimulated by TEX19. Endogenously expressed mL1-ORF1p in mouse testes represents a collection of protein molecules expressed from hundreds of variant copies of L1 at different genomic loci, therefore to allow us to correlate the abundance of L1-ORF1p with its encoding RNA more accurately, and to detect transient polyubiquitylated intermediates that are destined for proteasome-dependent degradation, we expressed engineered epitope-tagged hL1-ORF1p constructs in HEK293T cells. HEK293T cells do not endogenously express detectable levels of TEX19 (Reichmann et al. 2017) and *in vivo* ubiquitylation assays show that there is basal ubiquitylation of hL1-ORF1p, detectable as a ladder of hL1-ORF1p species in his_6_-myc-Ub pull-downs, in these cells (Figure 3A). The increasing molecular weights of these bands presumably correspond to increasing ubiquitylation of hL1-ORF1p. Furthermore, treating these cells with the proteasome inhibitor MG132 showed that hL1-ORF1p abundance is negatively regulated by the proteasome in the absence of TEX19 (Figure 3B). Interestingly, co-expression of *TEX19* during the *in vivo* ubiquitylation assay increasespolyubiquitylation of hL1-ORF1p (Figure 3C). *TEX19* increases the proportion of hL1-ORF1p-T7 that has at least four ubiquitin monomers, the minimum length of polyubiquitin chain required to target proteins to the proteasome (Thrower et al. 2000). Furthermore, expression of *TEX19* in these cells also reduces the abundance of hL1-ORF1p protein without any change in the abundance of its encoding RNA (Figure 3D). These gain-of-function data for *TEX19* mirror the loss-of-function data obtained from *Tex19.1*^*−/−*^ testes, confirm that the increased mL1-ORF1p levels in *Tex19.1* ^*−/−*^ testes are not a consequence of altered progression of *Tex19.1* ^*−/−*^ spermatocytes through meiosis (Crichton et al. 2017b; Öllinger et al. 2008), and suggest that *Tex19.1* orthologs function to post-translationally regulate L1-ORF1p abundance. The ubiquitylation and interaction data together suggests that, TEX19 orthologs regulate L1-ORF1p abundance through binding to L1-ORF1p and stimulating its polyubiquitylation and proteasome-dependent degradation.

**Figure 3.**
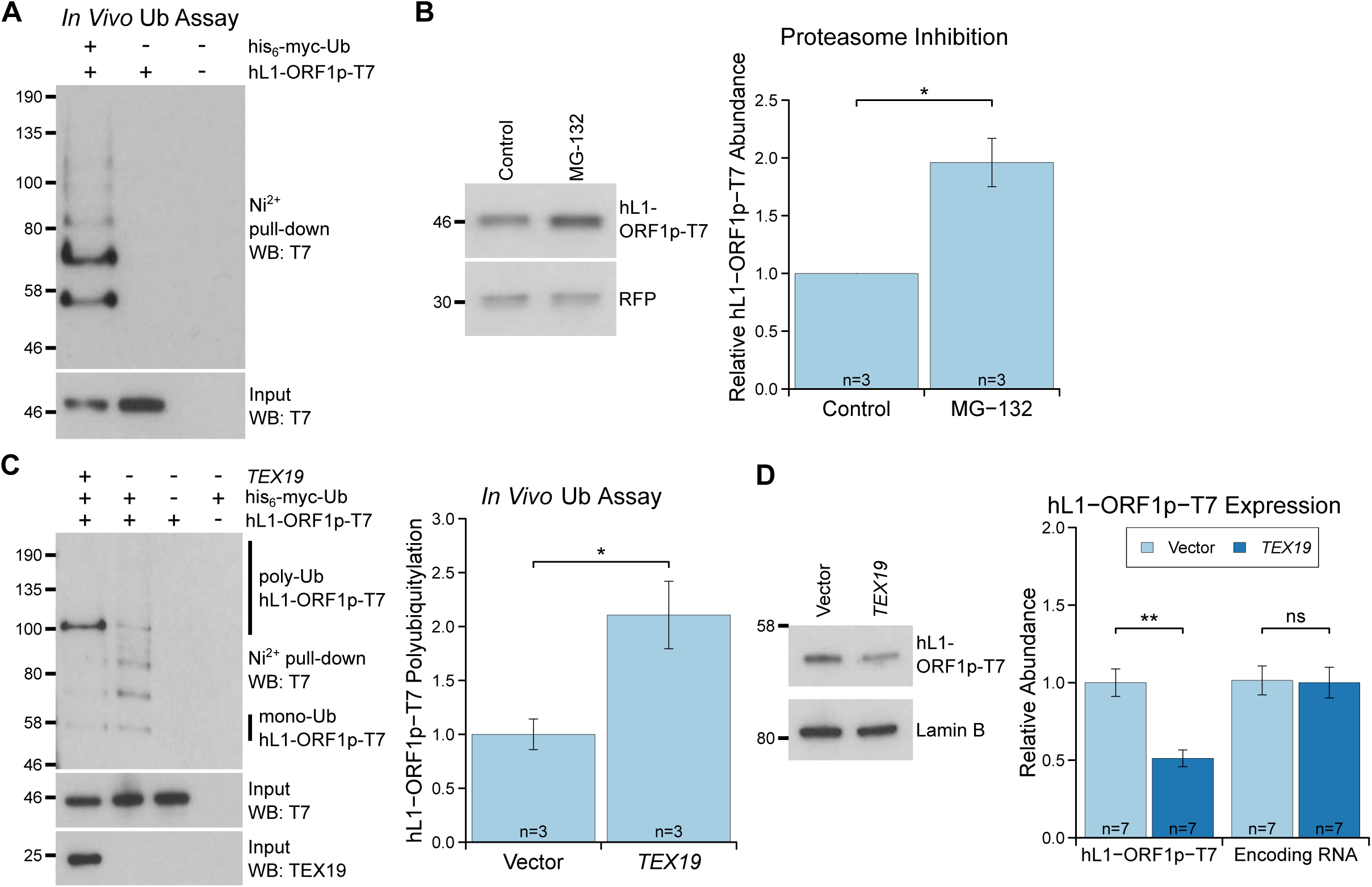
***TEX19* Stimulates Polyubiquitylation of hL1-ORF1p. A.***In vivo* ubiquitylation assay(Ub assay) for T7 epitope-tagged hL1-ORF1p in HEK293T cells. HEK293T cells were transfected with hL1-ORF1p-T7 and his_6_-myc-ubiquitin (his_6_-myc-Ub), and his_6_-tagged proteins isolated using Ni^2+^ agarose. Inputs and Ni^2+^ pull-downs were analysed by Western blotting for T7. **B.** Western blots and quantification of hL1-ORF1p-T7 abundance in HEK293T cells after treatment with either the proteasome inhibitor MG132 (50 µM, 7 hours) or DMSO as a vehicle control. HEK293T cells were co-transfected with hL1-ORF1p-T7 and RFP to control for transfection efficiency, and hL1-ORF1p-T7 abundance measured relative to RFP, then normalised to the DMSO controls for three independent transfections. MG132 treatment increases hL1-ORF1p-T7 abundance 1.96±0.21 fold. * p<0.05 (*t*-test). **C.**
*In vivo* ubiquitylation assay (Ub assay) for hL1-ORF1p-T7 in HEK293T cells in the presence and absence of human *TEX19*. Ni^2+^-pull downs were Western blotted (WB) with anti-T7 antibodies. Polyubiquitylated hL1-ORF1p-T7 containing four or more ubiquitin molecules (~100 kD band and above) was quantified relative to monoubiquitylated hL1-ORF1p-T7 (~58 kD band) and normalised to empty vector controls. Means ± SEM (1±0.14 and 2.11±0.31 for vector control and *TEX19* respectively) are indicated; * p<0.05 (*t*-test). **D.** Western blots of HEK293 FlpIn cells stably expressing hL1-ORF1p-T7 transfected with human *TEX19* or empty vector. Abundance of hL1-ORF1p-T7 protein and its encoding RNA were measured relative to lamin B and GAPDH respectively, and normalised to empty vector controls. Means ± SEM (1±0.09 and 0.51±0.06 for protein abundance and 1.01±0.09 and 1±0.10 for RNA abundance for vector control and *TEX19* respectively) are indicated; ** p < 0.01; ns indicates not significant (*t*-test); MW markers (kD) are indicated beside blots.

### *Tex19.1* Orthologs Restrict Mobilisation of L1 Reporters

L1-ORF1p has essential roles in L1 retrotransposition (Beck et al. 2011; Richardson et al. 2014a; Hancks and Kazazian 2016), therefore since *TEX19* orthologs bind to L1-ORF1p and negatively regulate its abundance, we investigated whether *Tex19.1* might inhibit L1 mobilisation in cultured cells. Engineered L1 retrotransposition assays with an EGFP retrotransposition reporter in HEK293T cells (Figure 4A) were used to measure the effect of *Tex19.1* on mobilisation of mouse L1 reporters. Expression of *Tex19.1* reduced the ability of both synthetic mouse L1 and a G_f_ type mouse L1 reporter to mobilise in these cells, but had no detectable effect on negative control reporters carrying inactivating mutations in ORF2 (Figure 4B). *Tex19.1* did not affect mobilisation of L1 retrotransposition reporters as potently as the L1 restriction factor *APOBEC3A* (Bogerd et al. 2006b), but still reduced L1 mobilisation by around 50% in this assay (Figure 4B). Mouse *Tex19.1* also restricts mobilisation of engineered human L1 reporters (Supplementary Figure S4A) althoughless efficiently than it restricts mouse L1 reporters. These data show that *Tex19.1* can function as a restriction factor for L1 mobilisation.

**Figure 4.**
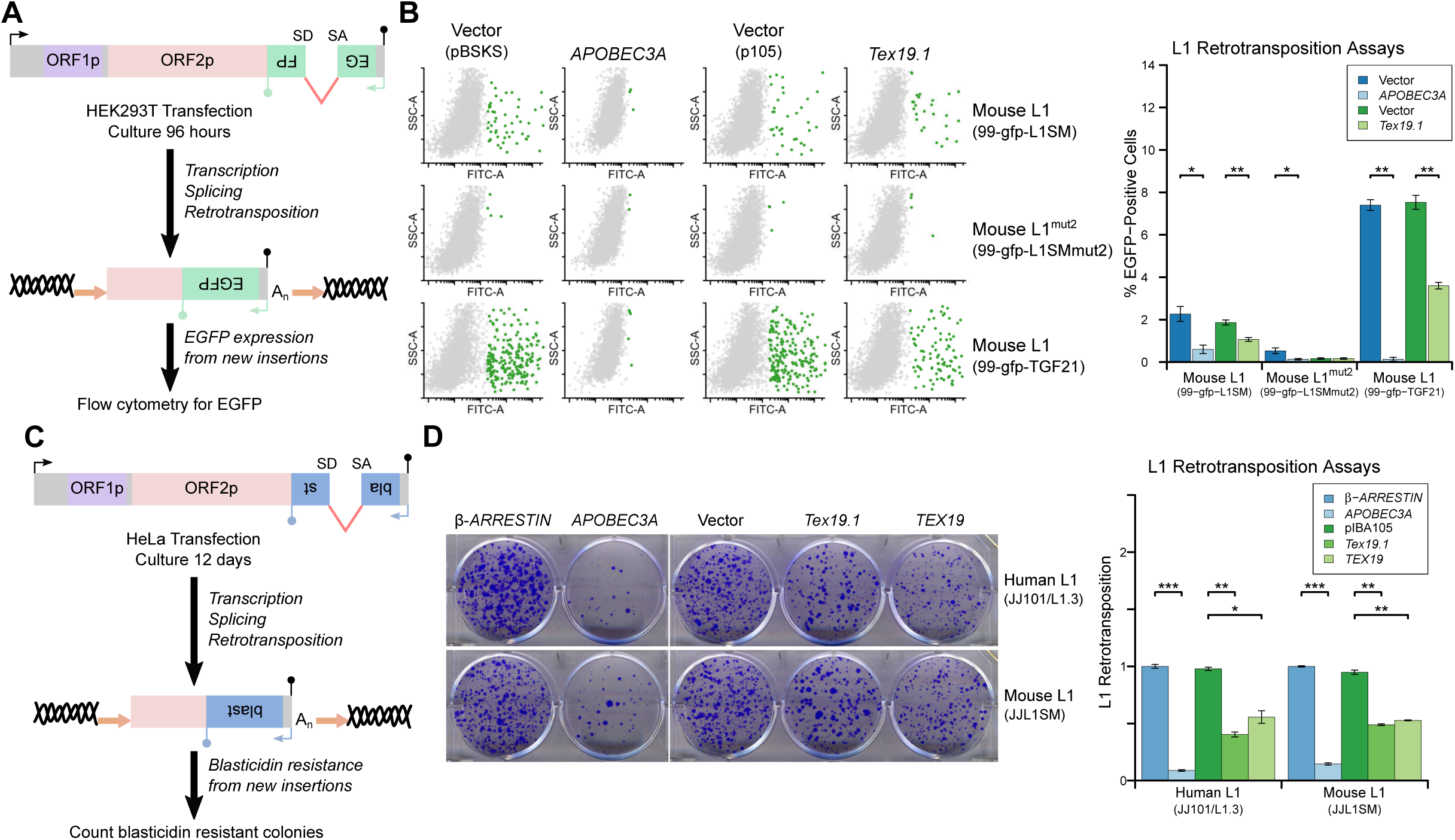
***TEX19* Orthologs Restrict L1 Mobilisation. A.** Schematic of engineered L1retrotransposition assay in HEK293T cells using EGFP as a reporter. **B.** Flow cytometry profiles from engineered mouse L1 retrotransposition assays performed as shown in panel A. HEK293T cells were co-transfected with a engineered mouse L1 retrotransposition constructs containing EGFP reporters (99-gfp-L1SM, 99-gfp-L1SMmut2, 99-gfp-TGF21), and either Strep-tagged *Tex19.1*, *APOBEC3A* (positive control) or empty vectors (pBSKS for *APOBEC3A*, pIBA105 for *Tex19.1*). EGFP fluorescence is plotted on the x-axis and side scatter on the y-axis of the flow cytometry profiles, and cells classed as EGFP-positive are shown in green. 99-gfp-L1SMmut2 carries missense mutations in the endonuclease and reverse transcriptase domains of ORF2p. * p<0.05; ** p<0.01 (*t*-test). **C.** Schematic of engineered L1 retrotransposition assays in HeLa cells using a blasticidin resistance reporter. **D.** Plates stained with 0.1% crystal violet showing blasticidin-resistant colonies from engineered L1 retrotransposition assays performed as shown in panel C. Human (JJ101/L1.3) and mouse (JJL1SM) L1 retrotransposition constructs containing blasticidin resistance reporters were co-transfected with β*-ARRESTIN* or *APOBEC3A* as negative and positive controls respectively, or with Strep-tagged mouse *Tex19.1*, Strep-tagged human *TEX19* or pIBA105 empty vector. Quantification of L1 retrotransposition was calculated relative to the β*-ARRESTIN* control. * p<0.05; ** p<0.01 (*t*-test).

Mouse *Tex19.1* expression is activated in response to DNA hypomethylation in multiple contexts (Hackett et al. 2012), and in humans *TEX19* is a cancer testis antigen expressed in multiple types of cancer (Feichtinger et al. 2012). We therefore tested whether expression of *TEX19* orthologs might be sufficient to restrict L1 mobilisation in multiple host cell types. L1 retrotransposition assays using blasticidin resistance reporters in HeLa cells (Figure 4C) showed that mouse *Tex19.1* similarly restricts mobilisation of a mouse and human L1 reporter by ~50% in this epithelial carcinoma cell line (Figure 4D). Human *TEX19* also restricts mobilisation of mouse and human L1 reporters by ~50% in these cells (Figure 4D). Similar effects on mobilisation of L1 reporters were also observed in U2OS osteosarcoma cells (Supplementary Figure S4B, Supplementary Figure S4C). Thus, *TEX19* orthologs are host restriction factors for L1 retrotransposition in mice and humans. Importantly, although we have also shown that TEX19 orthologs promote polyubiquitylation and degradation of L1-ORF1p, since TEX19 can directly bind to L1-ORF1p it is possible that this interaction also disrupts aspects L1-ORF1p function and contributes to TEX19-dependent restriction of L1 mobilisation. Moreover, there could be additional aspects of TEX19 function that may also be contributing to its ability to restrict L1 mobilisation. Indeed, it is not uncommon for host restriction factors to influence multiple aspects of retrotransposon or retroviral life cycles (Wang et al. 2010; Burdick et al. 2010; Goodier et al. 2012; Holmes et al. 2007).

### UBR2 Interacts With L1-ORF1p And Regulates L1 Independently Of *Tex19.1*Orthologs

The stoichiometric abundance of TEX19.1 and UBR2 in co-IPs in combination with the co-fractionation of all detectable TEX19.1 protein with UBR2 (Figure 2A, Figure 2C) suggests that *TEX19*-dependent polyubiquitylation of L1-ORF1p, and possibly also TEX19-dependent restriction of L1 mobilisation, is likely mediated by a TEX19.1-UBR2 complex. In contrast to *Tex19.1*, *Ubr2* is ubiquitously expressed (Supplementary Figure S5A) and UBR2 could contribute to basal ubiquitylation of L1-ORF1p in HEK293T cells (Figure 3A) and other somatic cell types. TEX19.1 could then simply stimulate this activity when transcriptionally activated by programmed DNA hypomethylation in the developing germline. A simple test of this model would be that TEX19.1-dependent effects on L1-ORF1p abundance or L1 mobilisation ought to be abolished in a *Ubr2* mutant background. However, the requirement for UBR2 to stabilise TEX19.1 protein (Yang et al. 2010) confounds analysis of the downstream requirement of UBR2 catalytic activity in TEX19.1-dependent functions: as TEX19.1 protein is unstable and undetectable in the absence of UBR2 (Yang et al. 2010), TEX19.1 might be expected to be unable to stimulate L1-ORF1p degradation or restrict L1 mobilisation regardless of whether the E3 ubiquitin ligase activity of UBR2 is required for these functions or not. Indeed, *Ubr2*^*−/−*^ testes largely phenocopy *Tex19.1* ^*−/−*^ testes, including transcriptional de-repression of MMERVK10C LTR retrotransposons (Crichton et al. 2017a).

To dissociate the effects of UBR2 on stability of TEX19.1 protein from potential effects on L1-ORF1p abundance and L1 mobilisation we therefore tested whether UBR2 can regulate L1 in the absence of effects on TEX19 stability by using somatic HEK293T cells. Interestingly, mouse UBR2 co-IPs with mL1-ORF1p in HEK293T cells (Figure 5A), a cell type that does not express any detectable TEX19(Reichmann et al. 2017), suggesting that UBR2 is able to regulate L1-ORF1p independently of any effects on TEX19 protein stability. UBR2 also interacts with mL1-ORF1^RA^pmutants that have reduced binding to RNA (Figure 5B), suggesting that this physical interaction is not mediated by L1 or any other RNA. Furthermore, these interactions are conserved in human L1-ORF1p (Supplementary Figure S5B, Supplementary Figure S5C). In addition, overexpression of UBR2 alone restricts mobilisation of a human L1 retrotransposition reporter (Figure 5C). Thus, at least when it is overexpressed, UBR2 is able to physically interact with L1-ORF1p and restrict mobilisation of L1 reporters.

**Figure 5.**
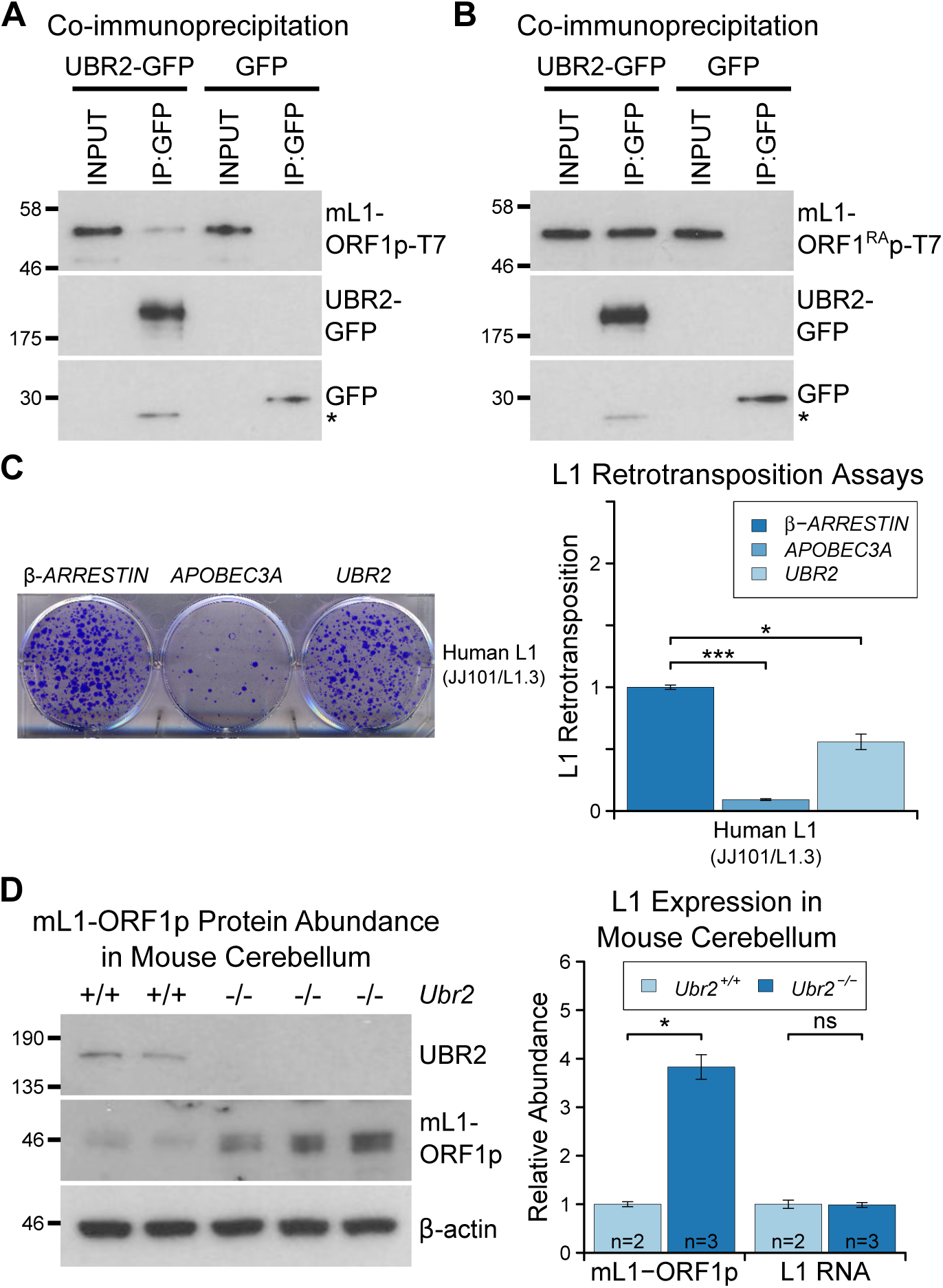
**The TEX19.1-Interacting Protein UBR2 Negatively Regulates mL1-ORF1p Abundance and L1 Mobilisation. A.** Co-immunoprecipitations (co-IPs) from HEK293T cells co-transfected with mL1-ORF1p-T7 and either mouse UBR2-GFP or GFP. IP inputs and IPs were Western blotted with T7 and GFP antibodies. A presumed cleavage product of UBR2-GFP running smaller than GFP itself is indicated with an asterisk. **C.** Plates from an engineered L1retrotransposition assay as described in Figure 4C stained with 0.1% crystal violet showing blasticidin-resistant colonies. Human (JJ101/L1.3) L1 retrotransposition construct was co-transfected with β*-ARRESTIN* or *APOBEC3A* as negative and positive controls respectively, or with UBR2-Flag. * p<0.05; *** p<0.01 (*t*-test). **D.** Western blots of endogenous UBR2 and mL1-ORF1p in P16 *Ubr2*^*+/+*^ and *Ubr2* ^*−/−*^ mouse cerebellum. β-actin was used as a loading control. Quantification of mL1-ORF1p-T7 and L1 mRNA relative to β-actin and normalised to *Ubr2* ^*+/+*^ control mice is also shown. Means ± SEM are indicated (1±0.05 and 3.82±0.25 for *Ubr2* ^*+/+*^ and *Ubr2* ^*−/−*^ respectively) * p<0.05; ns indicates not significant (*t*-test); MW markers (kD) are shown beside blots.

To test whether endogenously expressed UBR2 might regulate hL1-ORF1p abundance we generated *UBR2* mutant HEK293T cell lines by CRISPR/Cas9-mediated genome editing. However, these cell lines grew slowly and poorly in culture, presumably reflecting the normal cellular roles of UBR2 in cohesin regulation, DNA repair, and chromosome stability (Ouyang et al. 2006; Reichmann et al. 2017). Therefore, to allow a meaningful analysis of the role of endogenous UBR2 in L1 regulation we analysed *Ubr2* ^*−/−*^ mice (Supplementary Figure S5D, Supplementary Figure SE) which, despite having defects in spermatogenesis and female lethality, are otherwise grossly normal (Kwon et al. 2003). mL1, but not *Tex19.1*, is expressed in the brain (Wang et al. 2001; Muotri et al. 2010), therefore we used this tissue to assess whether *Ubr2* might have a *Tex19.1*-independent role in regulating mL1-ORF1p. Consistent with the physical interaction between UBR2 and mL1-ORF1p (Figure 5A), we found that mL1-ORF1p abundance is post-transcriptionally elevated in the cerebellum of *Ubr2* ^*−/−*^ mice (Figure 5D), suggesting that UBR2 may directly regulate polyubiquitylation and subsequent degradation of mL1-ORF1p *in vivo*. Interestingly, loss of *Ubr2* has no detectable effect on mL1-ORF1p abundance in the cerebrum (Supplementary Figure S4E), which may reflect cell type specific differences in L1 regulation or genetic redundancy between UBR-domain proteins (Tasaki et al. 2005). Nevertheless, regardless of this additional complexity in the cerebrum, the increased abundance of mL1-ORF1p in *Ubr2* ^*−/−*^ cerebellum demonstrates that endogenous *Ubr2* plays a *Tex19.1*-independent role in regulating mL1-ORF1p abundance *in vivo*.

*Ubr2* has numerous endogenous cellular substrates and host functions beyond regulating mL1-ORF1p (Ouyang et al. 2006; Reichmann et al. 2017; Sriram et al. 2011), but expression of *Tex19.1* in the germline or in response to DNA hypomethylation appears to stimulate a pre-existing activity of UBR2 to regulate mL1-ORF1p, possibly at the expense of UBR2’s activity towards some endogenous cellular substrates (Reichmann et al. 2017).

### *Tex19.1* Regulates mL1-ORF1p Abundance and Restricts L1 Mobilisation in Pluripotent Cells

As outlined earlier, L1 mobilisation is thought to occur primarily in pluripotent cells within the germline cycle (Kano et al. 2009), and regulation of L1 expression and mobilisation in these cells is likely to significantly impact on the ability of L1 to influence germline mutation and genome evolution. Therefore, we tested whether *Tex19.1*, which is expressed in pluripotent cells (Kuntz et al. 2008), has a role in regulating L1 expression and restricting L1 mobilisation in this cell type. We first investigated whether *Tex19.1* regulates mL1-ORF1p abundance in pluripotent mouse ESCs. Biochemical isolation of polyubiquitylated proteins and treatment with proteasome inhibitor suggests that endogenous mL1-ORF1p is polyubiquitylated and regulated by the proteasome in pluripotent mouse ESCs (Figure 6A, Figure 6B). hL1-ORF1p abundance is similarly regulated by the proteasome in human ESCs and human embryonal carcinoma cells (Supplementary Figure S6). In contrast to a previous report assessing the abundance of retrotransposon RNAs in ESCs derived from heterozygous mouse crosses (Tarabay et al. 2013), *Tex19.1*
^*−/−*^ mouse ESCs generated by sequential gene targeting (Supplementary Figure S7) in a defined genetic background and analysed at low passage number do not de-repress L1 RNA (Figure 6C). These *Tex19.1* ^*−/−*^ mouse ESCs contain elevated levels of endogenous mL1-ORF1p, but this increase in mL1-ORF1p levels is not accompanied by increased endogenous L1 mRNA levels (Figure 6C). Moreover, loss of *Tex19.1* does not affect transcription or translation of L1 reporter constructs in ESCs (Supplementary Figure S8). Thus, like in male germ cells (Figure 1), *Tex19.1* functions to post-translationally repress mL1-ORF1p in pluripotent cells.

**Figure 6.**
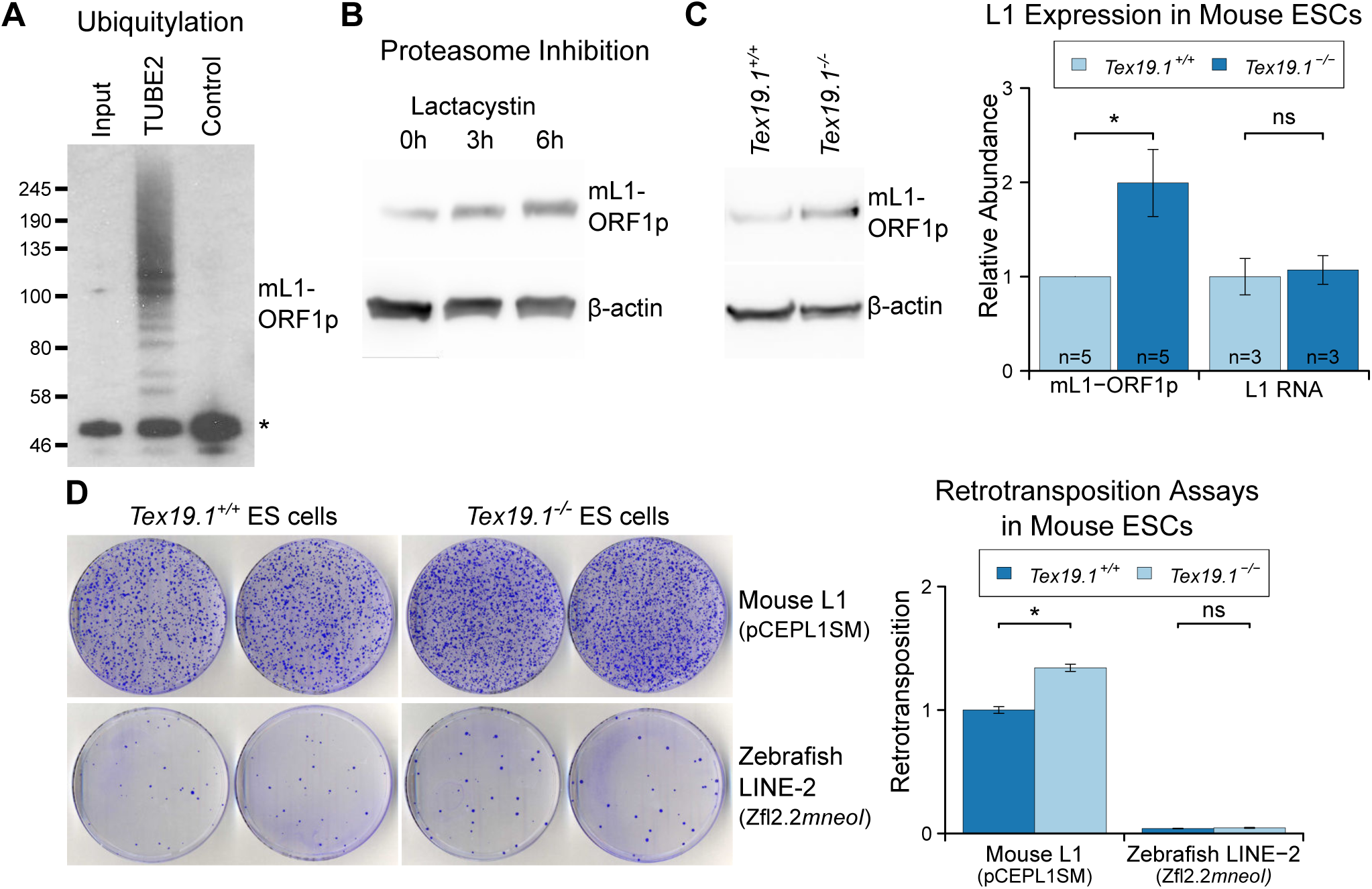
***Tex19.1* Negatively Regulates mL1-ORF1p Abundance And L1 Mobilisation In Mouse ESCs. A.** Mouse ESC lysates (input) were incubated with polyubiquitin-binding TUBE2beads or control agarose beads and Western blotted for endogenous mL1-ORF1p. Non-specific binding of non-ubiquitylated mL1-ORF1p is detectable (asterisk), in addition to specific enrichment of polyubiquitylated mL1-ORF1p with TUBE2. **B.** Western blot for endogenous mL1-ORF1p after treatment with 25 µM lactacystin proteasome inhibitor for the indicated times. β-actin is a loading control. **C.** Western blot for endogenous mL1-ORF1p in *Tex19.1*
^*+/+*^ and *Tex19.1* ^*−/−*^ mouse ESCs. mL1-ORF1p abundance (Western blot) and L1 RNA abundance (qRT-PCR) were quantified relative to β-actin and normalised to *Tex19.1* ^*+/+*^ ESCs. Means ± SEM are indicated (1±0 and 1.99±0.36 for protein and 1±0.19 and 1.07±0.15 for RNA for *Tex19.1* ^*+/+*^ and *Tex19.1* ^*−/−*^ respectively); * p<0.05; ns indicates not significant (*t*-test). **D.** Neomycin-resistant colonies from L1 retrotransposition assays in *Tex19.1* ^*+/+*^ and *Tex19.1* ^*−/−*^ ESCs. ESCs were transfected with LINE retrotransposition constructs carrying the *mneoI* reporter cassette and either synthetic mouse L1 (pCEPL1SM) or zebrafish LINE-2 (Zfl2.2) sequences, the number of neomycin-resistant colonies counted, and retrotransposition frequency calculated relative to *Tex19.1* ^*+/+*^ ESCs transfected with pCEPL1SM. * p<0.05; ns indicates not significant (*t*-test); error bars indicate SEM.

We next tested whether loss of *Tex19.1* also results in increased mobilisation of mouse L1 reporters in pluripotent ESCs. Although L1 retrotransposition assays have previously been performed in pluripotent human cells (Wissing et al. 2011; Garcia-Perez et al. 2007, 2010; Wang et al. 2014; Klawitter et al. 2016), this assay had not been adapted to mouse ESCs and, to our knowledge, no restriction factor has yet been shown to restrict mobilisation of L1 reporters in mouse pluripotent cells or germ cells. Adaptation of the L1 retrotransposition assay using a blasticidin resistance retrotransposition reporter (Figure 4C) to mouse ESCs resulted in the appearance of blasticidin resistant colonies that could be suppressed by either introducing the N21A mutation (Alisch et al. 2006) into the endonuclease domain of ORF2, or co-transfection of the L1 restriction factor APOBEC3A (Bogerd et al. 2006b) (Supplementary Figure S9). Thus, the adapted L1 retrotransposition assay appears to reflect *bone fide* mobilisation of L1 reporters. Interestingly, mobilisation of L1 reporter constructs is elevated around 1.5-fold in *Tex19.1*^*−/−*^ ESCs (Figure 6D, Supplementary Figure S9). However, loss of *Tex19.1* does not influence the mobility of an engineered zebrafish LINE-2 reporter that lacks ORF1p (Kajikawa et al. 2012) (Figure 6D). Thus, these data suggest that the role of endogenously expressed *Tex19.1* in mouse pluripotent cells is to restrict L1 mobilisation, and thereby promote trans-generational genome stability.

## Discussion

This study identifies *Tex19.1* as a host restriction factor for L1 in the mammalian germline. We have previously reported that *Tex19.1* plays a role in regulating the abundance of retrotransposon RNAs (Öllinger et al. 2008; Reichmann et al. 2012, 2013), which appears to reflect transcriptional de-repression of specific retrotransposons (Crichton et al. 2017a). Although loss of *Tex19.1* results in de-repression of L1 RNA in placenta (Reichmann et al. 2013), L1 RNA abundance is not affected by loss of *Tex19.1* in male germ cells (Öllinger et al. 2008) or, in contrast to a previous report (Tarabay et al. 2013), in mouse ESCs (Figure 6). Indeed here we show that *Tex19.1* has a role in post-translational regulation of L1-ORF1p steady-state levels in these cells. Thus, *Tex19.1* appears to regulate retrotransposons at multiple stages of their life cycle. It is possible that *Tex19.1* is affecting different E3 ubiquitin ligases, or different E3 ubiquitin ligase substrates, in order to repress different stages of the retrotransposon life cycle. However, loss of *Tex19.1* results in a 1.5-fold increase in mobilisation of L1 reporters in pluripotent cells. Since L1 mobilisation mostly takes place in the pluripotent phase of the germline cycle, and new L1-dependent mobilisation events are thought to be inherited by one in every twenty human births (Kazazian 1999), *TEX19* activity could potentially be preventing new retrotransposition events from being inherited by up to 3 million births annually. Retrotransposons appear to provide functions that are advantageous for mammalian development and evolution (Garcia-Perez et al. 2016), and the activity of restriction mechanisms like the TEX19-dependent mechanism we have described here, that control the ability of retrotransposons to mobilise, rather than eliminate their transcriptional activity altogether, could potentially allow retrotransposons to participate in and drive the evolution of key gene regulatory networks in pluripotent cells while minimising their mutational load on the germline genome.

Our data suggests that L1-ORF1p is post-translationally modified by ubiquitylation in somatic and germline cells. Phosphorylation of L1-ORF1p has been previously reported in somatic tissues and is required for L1 retrotransposition in these cells (Cook et al. 2015). However, we are not aware of any previous reports that post-translational modifications of L1-ORF1p are present in the germline, particularly in the pluripotent phase of the germline cycle when L1 retrotransposition is thought to primarily occur (Kano et al. 2009). Post-translational regulation of L1 potentially provides an additional layer of genome defence that could be particularly important during periods of epigenetic reprogramming in early embryogenesis or in the developing primordial germ cells when transcriptional repression of retrotransposons might be more relaxed (Molaro et al. 2014; Fadloun et al. 2013). Indeed, the sensitivity of *Tex19.1* expression to DNA hypomethylation (Hackett et al. 2012) will allow post-translational suppression of L1 to be enhanced during these stages of development. Post-translational regulation of L1 is also likely important to limit the activity of L1 variants that evolve to escape transcriptional repression and will provide a layer of genome defence while the host adapts its KRAB zinc-finger protein repertoire to these new variants (Jacobs et al. 2014). Analysis of L1 evolution shows that regions within L1-ORF1p are under strong positive selection suggesting that host restriction systems are targeting L1-ORF1p post-translationally and impacting on evolution of these elements (Boissinot and Furano 2001; Sookdeo et al. 2013). Although this evidence for post-translational restriction factors acting on L1-ORF1p has been known for over 15 years, to our knowledge no host factors have been identified that directly bind to L1-ORF1p and restrict L1 mobilisation. It is possible that the physical interactions between L1-ORF1p and TEX19: UBR2 that we describe here are contributing to these selection pressures acting on L1-ORF1p. While UBR2 is able to target L1-ORF1p in the absence of TEX19, evolution of a less constrained TEX19 adapter to provide a further link between UBR2 and L1-ORF1p could potentially resolve the contradictory pressures on UBR2 to maintain interactions with some endogenous cellular substrates while targeting a rapidly evolving retrotransposon protein for degradation.

Our data indicates that TEX19.1 likely exists in a complex with UBR2 in ESCs, and that TEX19.1 stimulates a basal activity of UBR2 to bind to and polyubiquitylate L1-ORF1p (Figure 7). Ubr1, a yeast ortholog of UBR2, has different binding sites for different types of substrate (Xia et al. 2008). Ubr1 can bind to proteins that have specific residues at their N-termini (N-end rule degrons), and to proteins that have more poorly defined non-N-terminal internal degrons. Full-length human L1-ORF1p does not have a potential N-end rule degron at its N-terminus (Kim et al. 2014), and the interaction between UBR2 and L1-ORF1p likely reflects an internal degron in the retrotransposon protein. One of the known internal degron substrates of yeast Ubr1 is CUP9, a transcription factor that regulates expression of a peptide transporter (Turner et al. 2000). Binding of Ubr1 to the CUP9 internal degron is activated by specific dipeptides binding to the N-end rule degron binding sites, which inhibits binding of N-end rule degrons, allosterically relieves the activity of an autoinhibitory domain within Ubr1, stimulates binding to the CUP9 internal degron, and increase processivity of Ubr1 resulting in increased polyubiquitylation of CUP9 (Du et al. 2002; Xia et al. 2008; Turner et al. 2000). This generates a positive feedback loop to activate transcription of a peptide transporter when these peptides are present. The effect of these dipeptides on Ubr1 activity in yeast strongly resonates with the effects of TEX19 orthologs on UBR2 activity in mammals. TEX19 binds to UBR2 and inhibits its activity towards N-end rule substrates (Reichmann et al. 2017), but stimulates polyubiquitylation of L1-ORF1p, possibly through an internal degron in L1-ORF1p. The direct interaction between TEX19.1 and L1-ORF1p could further enhance L1-ORF1p binding to UBR2 by stabilising the highly flexible L1-ORF1p trimers (Khazina et al. 2011) in a conformational state that exposes an internal degron and favours their ubiquitylation. Thus, TEX19 orthologs appear to function, at least in part, by re-targeting UBR2 away from N-end rule substrates and towards a retrotransposon substrate. However, the direct interaction between TEX19.1 and L1-ORF1p means that it is possible that TEX19.1 is interfering with L1-ORF1p function in multiple ways in order to restrict L1 mobilisation. Thus, while one outcome of this interaction appears to be increased polyubiquitylation and degradation of L1-ORF1p, the interaction between TEX19.1 and L1-ORF1p could also interfere with the nucleic acid chaperone activity of L1-ORF1p (Martin et al. 2005), or its interactions with either L1-encoded or host-encoded molecules. Further work is needed to determine whether these or other additional mechanisms are contributing to the ability of TEX19.1 to restrict L1 mobilisation.

**Figure 7.**
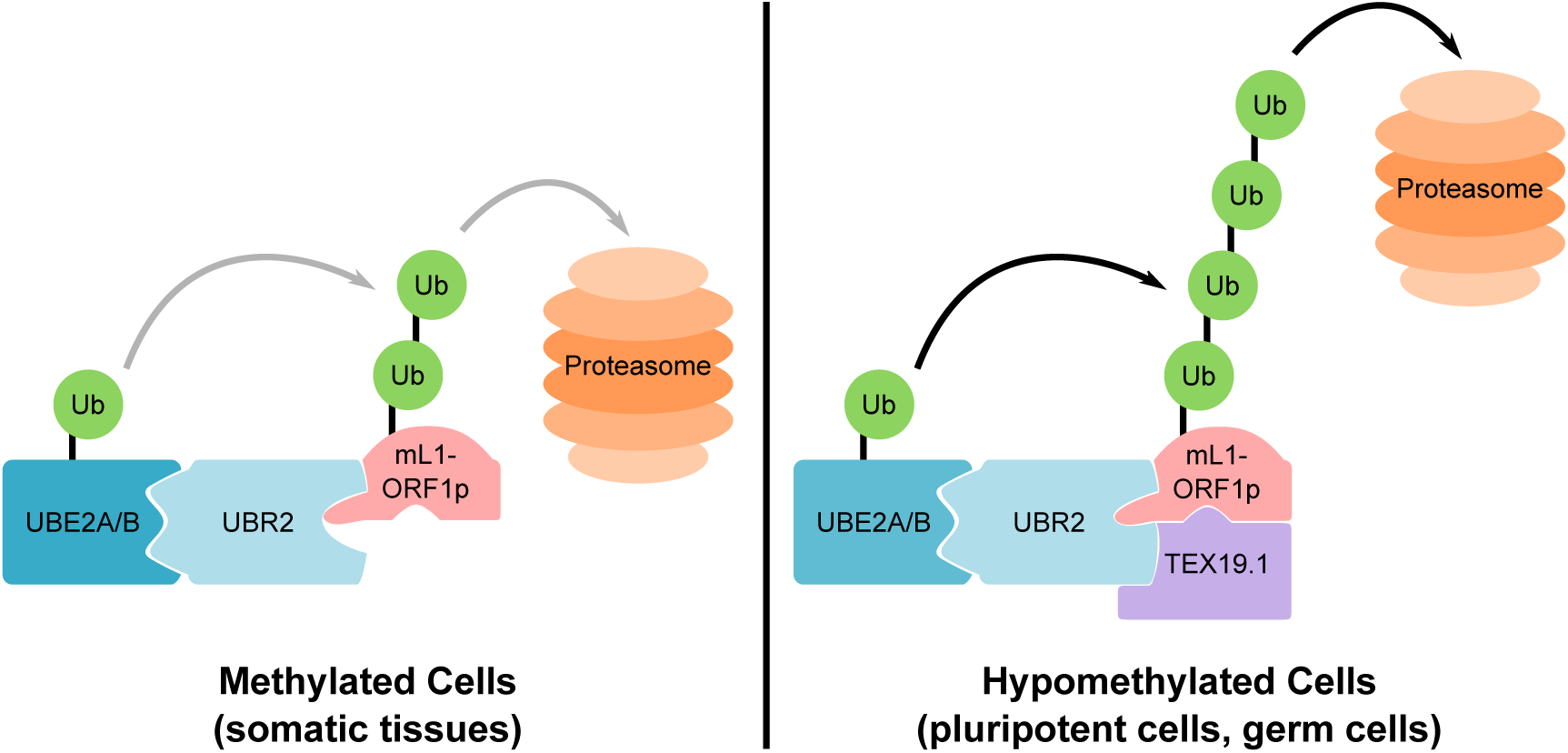
**Model For UBR2:TEX19.1-Mediated Polyubiquitylation of mL1-ORF1p**. In methylated somatic cells, the RING domain E3 ubiquitin ligase UBR2 and its cognate E2 ubiquitin conjugating enzyme UBE2A/B can interact with mL1-ORF1p and catalyse ubiquitylation and proteasome-dependent turnover of this protein. TEX19.1 in hypomethylated cells, including pluripotent cells and germ cells, interacts with both UBR2 and mL1-ORF1p, stimulating further polyubiquitylation and proteasome-dependent turnover of mL1-ORF1p. The interaction between TEX19.1 and UBR2 concomitantly inhibits the activity of UBR2 towards N-end rule substrates (Reichmann et al. 2017).

The data presented here suggests that programmed DNA hypomethylation in the mouse germline extends beyond activating components of the PIWI-piRNA pathway (Hackett et al. 2012) to include enhancing the activity of the ubiquitin-proteasome system towards retrotransposon substrates. Activation of post-translational genome-defence mechanisms may allow mammalian germ cells to safely transcribe retrotransposons by preventing these transcripts from generating RNPs that can mutate the germline genome (Supplementary Figure S10). The retrotransposon transcripts can then be processed into piRNAs and used to identify retrotransposon loci where epigenetic silencing needs to be established. *De novo* establishment of epigenetic silencing at retrotransposons in the *Arabidopsis* germline involves transfer of small RNAs between a hypomethylated vegetative cell and a germ cell (Slotkin et al. 2009), whereas these processes happen sequentially in the same germ cell in mammals (Supplementary Figure S10). Therefore the ability to enhance post-translational control of retrotransposons may be a key feature of epigenetic reprogramming in the mammalian germline that limits the trans-generational genomic instability caused by retrotransposon mobilisation.

## Acknowledgments

We thank the MRC (intramural programme grants to I. R. A (MC_PC_U127580973) and to R. R. M. (MC_PC_U127574433); doctoral training awards to J. R. and C. J. P) for funding. J. L. G. P.’s lab is supported by CICE-FEDER-P12-CTS-2256, Plan Nacional de I+D+I 2008-2011 and 2013-2016 (FIS-FEDER-PI11/01489 and FIS-FEDER-PI14/02152), PCIN-2014-115-ERA-NET NEURON II, the European Research Council (ERC-Consolidator ERC-STG-2012-233764), by an International Early Career Scientist grant from the Howard Hughes Medical Institute (IECS-55007420) and by The Wellcome Trust-University of Edinburgh Institutional Strategic Support Fund (ISFF2). E. K and O. W. are supported by the Max Planck Society. R. E. was funded by the Norwegian Cancer Society (2293664-2011) and Department of Biosciences, University of Oslo. We thank Sandy Martin (University of Colorado), Alex Bortvin (Carnegie Institute, Baltimore), John V. Moran (Universiy of Michigan), Jef D. Boeke (New York University), John Goodier (Johns Hopkins University, Baltimore), Norihiro Okada (National Cheng Kung University), Niki Gray, Matt Brook, Sara Heras, and Mark Ditzel (all University of Edinburgh) for generously sharing reagents and technical expertise, the flow cytometry, imaging and animal facilities for technical support, and Wendy Bickmore and Javier Caceres (both University of Edinburgh) for critical comments on the manuscript.

### Author Contributions

M. M. performed the ubiquitylation assays, co-IPs, and HEK293T, ES cell and *Ubr2* ^*−/−*^ mouse tissue analysis; M. G-C, performed ES cell analysis, RTRN assays and confocal microscopy; J. R. analyzed L1 expression and translation in *Tex19.1* ^*−/−*^ mouse testes; E. Z. and O. W. conceived, designed, performed and analysed experiments testing TEX19: L1-ORF1p interactions with bacterially-expressed proteins; C. S-P., P. P. and L. S. performed RTRN assays; A. R. M. generated *Tex19.1* ^*−/−*^ ES cells; C. J. P. peformed qRT-PCR assays on *Ubr2* ^*−/−*^ mouse tissues, D. R. performed mouse husbandry, genotyping and tissue collection; C.-C. H. generated *Tex19.1*-expressing stable cell lines, R. E. and J. R. performed the gel filtration analysis; R. R. M., J. L. G.-P. and I. R. A. conceived and designed the study. J. L. G.-P. and I. R. A. wrote the paper. M. M. and M. G.-C. contributed equally to the study. All authors discussed the results and commented on the manuscript.

### Materials and Methods

#### Mice

*TEX19.1* mutant mice on a C57BL/6 genetic background were maintained and genotyped as described (Öllinger et al. 2008; Reichmann et al. 2012). *Tex19.1* ^*+/−*^ heterozygotes have no detectable testis phenotype and indistinguishable sperm counts from wild-type animals (Öllinger et al. 2008), and prepubertal *Tex19.1* ^*−/−*^ homozygotes were typically compared with heterozygous littermates to control for variation between litters. *Ubr2* ^*−/−*^ mice were generated by CRISPR/Cas9 double nickase-mediated genome editing in zygotes (Ran et al. 2013). Complementary oligonucleotides (Supplementary Table 2) targeting exon 3 of UBR2 were annealed and cloned into plasmid pX335 (Cong et al. 2013), amplified by PCR, then *in vitro* transcribed using a T7 Quick High Yield RNA Synthesis kit (NEB) to generate paired guide RNAs. RNA encoding the Cas9 nickase mutant (50 ng/µl, Tebu-Bio), paired guide RNAs targeting exon 3 of UBR2 (each at 25 ng/µl), and 150 ng/µl single-stranded DNA oligonucleotide repair template (Supplementary Table 2) were microinjected into the cytoplasm of C57BL/6 × CBA F2 zygotes. The repair template introduces an *Xba*I restriction site and mutates cysteine-121 within the UBR domain of UBR2 (Uniprot Q6WKZ8-1) to a premature stop codon. The zygotes were then cultured overnight in KSOM (Millipore) and transferred into the oviduct of pseudopregnant recipient females. Pups were genotyped for the presence of the *Xba*I restriction site. The *Ubr2*^*−/−*^ male mice generated in this way have no overt phenotypes except testis defects and infertility and *Ubr2* ^*−/−*^ females are born at sub-Mendelian ratios, all as previously described for *Ubr2* ^*−/−*^ mice generated by gene targeting in ESCs (Kwon et al. 2003). The day of birth was designated P1, and mice were culled by cervical dislocation. Mouse experiments were performed in accordance with local ethical guidelines and under authority of UK Home Office Project Licence PPL 60/4424.

#### Cell Culture

We used cell lines that were previously shown to support retrotransposition of engineered L1 constructs or *Tex19.1*^*−/−*^ models generated in this study. Cell lines were maintained at 37°C in 5% CO_2_. HEK293T and U2OS cells were obtained from ATCC and HeLa cells were provided by John V. Moran (University of Michigan, US). These cell lines were grown in Dulbecco’s Modified Eagle’s Media (DMEM) supplemented with 10% foetal calf serum, 1% penicillin-streptomycin, and 1% L-glutamine. E14 ESCs were cultured on gelatinised flasks in 2i culture conditions (1:1 DMEM/F12 media: neurobasal media supplemented with N2 and B27, 10% foetal calf serum, 1% L-glutamine, 0.1% β-mercaptoethanol, 1 µM PD0325901 (StemMACS), and 3 µM CHIR99021 (StemMACS).

Hamster XR-1 cells (Stamato et al. 1983) were provided by Thomas D. Stamato (The Lankenau Institute fro Medical Research, US) and grown in DMEM low glucose medium containing 10% foetal calf serum, 1% L-glutamine, 1% penicillin-streptomycin and 0.1 mM non-essential amino acids. Human PA-1 cells (Zeuthen et al. 1980) were obtained from ATCC and grown in Minimal Essential Media (MEM) supplemented with 10% heat-inactivated foetal calf serum, 1% L-glutamine, 1% penicillin-streptomycin and 0.1 mM non-essential amino acids. H9 human ESCs (Thomson et al. 1998) were obtained from Wicell and cultured and passaged as previously described using conditional media (CM) (Garcia-Perez et al. 2007). To prepare CM, human foreskin fibroblasts (obtained from ATCC) were mitotically inactivated with 3000-3200 rads γ-irradiation, seeded at 3x10^6^ cells/225 cm^2^ flask and cultured with hESC media (KnockOut DMEM supplemented with 4 ng/ml bFGF, 20% KnockOut serum replacement, 1 mM L-glutamine, 0.1 mM β-mercaptoethanol and 0.1 mM non-essential amino acids) for at least 24 hours before media harvesting. We collected CM 24, 48 and 72 hours after seeding. H9 human ESCs (Wicell) were maintained on Matrigel (BD Biosciences)-coated plates in human foreskin fibroblast-conditionedmedia. The absence of *Mycoplasma* in cultured cells was confirmed once a month using a PCR-based assay (Minerva). Single tandem repeat genotyping was done at least once a year to ensure the identity of the cell lines used.

#### Generation Of Stable Cell Lines

ES and HEK293 cell lines stably expressing TEX19.1-YFP or YFP alone were generated by transfecting E14 ESCs or HEK293 cells with linearised pCAG-TEX19.1-YFP and pCAG-YFP expression plasmids (Supplementary Table 3) and selecting for the G418 resistance cassette. Stable cell lines were flow sorted to select for YFP expression. For pCAG-YFP transfection, the cell lines were flow sorted to select for cells expressing YFP at similar levels to the pCAG-TEX19.1-YFP cell lines. Stable Flp-In-293 cells (Invitrogen) expressing T7-tagged hL1-ORF1p from a CMV promoter at the Flp-In locus were generated using the pcDNA5⁄FRT Flp-In vector, and selected using 100 μg/ml hygromycin and 100 μg/ml Zeocin according to the supplier’s instructions. Validation of these cell lines is shown in Extended Data Figure 10.

#### Generation Of *Tex19.1*^*−/−*^ ESCs

*Tex19.1* ^*−/−*^ ESCs were generated by sequential targeting of E14 ESCs. The *Tex19.1* targeting vector was generated by inserting an IRES-GFP cassette into position chr11:121147942 (mm10 genome assembly) in the 3' untranslated region of *Tex19.1* in a bacterial artificial chromosome (BAC) by BAC recombineering (Liu et al. 2003). A 13 kb region (chr11:121143511-121156687) containing *Tex19.1* was gap-repaired into PL253 (Liu et al. 2003), then a LoxP site from PL452 was recombined upstream of the coding exon at position chr11:121146376, and an Frt-flanked neomycin-resistance cassette and second LoxP site from PL451 (Liu et al. 2003) recombined downstream of the coding exon at chr11:121148877. E14 ESCs were electroporated with the resulting targeting vector, selected for neomycin resistance, and correct integrants identified by PCR. The *Tex19.1* coding exon in the targeted allele was removed by transfection with a Cre-expressing plasmid, and the resulting cells electroporated with the targeting vector again, selected for neomycin resistance, and correct integrants on the second *Tex19.1* allele identified by PCR. ESCs were then transiently transfected with a Flp-expressing plasmid to generate a conditional *Tex19.1* ^*fl*^ allele. This was subsequently converted to a *Tex19.1* ^*−*^ allele by transient transfection with a Cre-expressing plasmid to remove the *Tex19.1* coding exon. ESCs were cultured in gelatinised flasks in LIF+ (Glasgow Modified Eagle’s Media, 10% foetal calf serum, 1% non-essential amino acid, 1% sodium pyruvate, 1% penicillin-streptomycin, 1% L-glutamine, 0.001% β-mercaptoethanol, and 0.2% leukaemia inhibitory factor-conditioned media) during the generation of *Tex19.1* ^*−/−*^ ESCs, then low passage *Tex19.1* ^*−/−*^ ESCs with a euploid karyotype were used for experiments after transitioning to 2i culture conditions for at least 14 days. Validation of this ESC line is shown in Extended Data Figure 10.

#### qRT-PCR

RNA was isolated from cells or tissues using TRIzol reagent (Life Technologies), treated with DNAse (DNAfree, Ambion) and used to generate random-primed cDNA (First Strand cDNA Kit, Life Technologies) as described by the suppliers. qPCR was performed on the cDNA using the SYBR Green PCR System (Stratagene) and a CFX96 Real-Time PCR Detection System (Bio-Rad). Control qRT-PCR reactions were performed in the absence of either reverse transcriptase or qPCR template to verify the specificity of any qRT-PCR signals obtained. Primers were validated to perform at >90% efficiency in the qRT-PCR assay, and expression quantified using the 2^−∆∆Ct^ method (Livak and Schmittgen 2001). Alternatively, qPCR was performed using SYBR Select Master Mix (Applied Biosystems) and a Light Cycler 480 II (Roche), and expression quantified using the relative standard curve method as described by the suppliers.

#### Western Blotting

Tissue or cells were homogenised in 2× Laemmli SDS sample buffer (Sigma) with a motorised pestle, then boiled for 2-5 minutes and insoluble material pelleted in a microcentrifuge. Protein samples were resolved on pre-cast Bis-Tris polyacrylamide gels in MOPS running buffer (Invitrogen), or Tris-Acetate polyacrylamide gels in Tris-Acetate SDS running buffer (Invitrogen) and Western blotted to PVDF membrane using a GENIE blotter (Idea Scientific) or the iBlot Transfer system (Invitrogen). Membranes were blocked with 5% non-fat skimmed milk powder in PBST (PBS, 0.1% Tween-20), then incubated with primary antibodies (Supplementary Table 4) diluted in blocking solution. Membranes were then washed with PBST and, if required, incubated with peroxidase-conjugated secondary antibody in blocking solution. Membranes were washed in PBST and bound secondary antibodies detected using West Pico Chemiluminescent Substrate (Thermo Scientific). Western blots were quantified using ImageJ (Schneider et al. 2012)

#### Immunostaining

Immunostaining on P16 testes was performed by fixing decapsulated P16 testes in 4% paraformaldehyde in PBS, embedding the tissue in paraffin wax, and cutting 6 μm sections on a microtome. Sections were de-waxed in xylene, rehydrated, and antigen retrieval was performed by boiling slides in a microwave for 15 mins in 10 mM sodium citrate pH 6. Sections were blocked in PBS containing 10% goat serum, 3% BSA, 0.1% Tween-20, then incubated in 1:300 rabbit anti-L1 ORF1p primary antibody diluted in blocking solution. Sections were then washed with PBS, incubated in 1:500 Alexa Fluor-conjugated secondary antibodies (Life Technologies), washed with PBS again, then mounted under a coverslip using Vectashield mounting media containing DAPI (Vector Laboratories). Slides were imaged on a Zeiss Axioplan II fluorescence microscope equipped with a Hamamatsu Orca CCD camera. Anti-mL1-ORF1p fluorescence intensity was measured per unit area, with slides immunostained with non-specific rabbit IgG and secondary antibodies used to calculate and subtract background.

#### Polysome Gradients

Polysome gradients were prepared as described (Gillian-Daniel et al. 1998). In brief, P18 testes were homogenised in 200 μl lysis buffer (20 mM HEPES pH 7.4, 150 mM KCl, 5 mM DTT, 5 mM MgCl_2_, 100 U/mL RNasein, Complete protease inhibitors (Roche), 10 nM calyculin A, 150 μg/Ml cycloheximide) then NP-40 added to 0.5% and the samples incubated on ice for 10 minutes. After centrifugation at 12,000*g* for 5 minutes at 4°C the soluble supernatant was layered onto an 11 mL 10-50% linear sucrose prepared in gradient buffer (20 mM HEPES pH 7.4, 250 mM KCl, 5 mM DTT, 10 mM MgCl_2_, 1 μg/μL heparin), then centrifuged in a SW41Ti rotor (Beckman) for 120 minutes at 38,000rpm at 4°C. 1 mL fractions were collected and absorbance of RNA at 254nm was recorded by using a UV monitor. To isolate RNA, fractions were digested with 20 μg/μL proteinase K in presence of 1% SDS and 10 mM EDTA for 30 minutes at 37°C then RNAs recovered using Trizol LS reagent (Invitrogen). To isolate proteins, fractions were precipitated with methanol/chloroform and pellets resuspended by boiling in Laemmli SDS sample buffer.

#### Oligo(dT) Pull-Downs

P16 testes were homogenised with a motorised pestle in lysis buffer (20 mM HEPES pH 7.4, 150 mM KCl, 5 mM DTT, 5 mM MgCl_2_) supplemented with 100 U/mL RNasein, Complete protease inhibitors (Roche) and insoluble debris removed by centrifugation (12,000*g*, 5 minutes at 4°C). Oligo(dT)-cellulose beads (Ambion) were blocked in lysis buffer containing 5% BSA for 1 h at 4°C, then incubated with lysate for 1 h at 4°C. Oligo(dT)-cellulose beads were washed three times with lysis buffer, and bound proteins eluted by boiling in Laemmli SDS sample buffer and analysed by Western blotting. For competition assays, 200 μg of a 25-mer poly(A) oligonucleotide (Sigma Genosys) was incubated with the oligo(dT)-cellulose beads for 30 min at 4°C before the addition of lysates.

#### Isolation Of TEX19.1-YFP Complexes

Cytoplasmic extracts were prepared as described (Wright et al. 2006). Briefly, stable YFP or TEX19.1-YFP ESCs growin in LIF+serum conditions were resuspended in 3 volumes buffer A (10 mM HEPES pH 7.6, 15 mM KCl, 2 mM MgCl_2_, 0.1 mM EDTA, 1 mM DTT, 0.2 mM PMSF, Complete protease inhibitors (Roche)) and incubated on ice for 30 mins. Cells were lysed in a Dounce homogeniser, one-tenth volume buffer B (50 mM HEPES pH 7.6, 1 M KCl, 30 mM MgCl_2_, 0.1 mM EDTA, 1% NP-40, 1 mM DTT, 0.2 mM PMSF) added, then the lysate centrifuged twice for 15 minutes at 3400*g* at 4°C to deplete nuclei. Glycerol was added to a final volume of 10%, the extracts centrifuged at 12,000*g* for 5 minutes at 4°C, pre-cleared with protein A agarose beads (Sigma) then with blocked agarose beads (Chromotek), before incubation with GFP-Trap agarose beads (Chromotek) for 90 minutes at 4°C. Beads were collected by centrifugation at 2700*g* for 2 minutes at 4°C, washed three times with 9:1 buffer A: buffer B, and protein eluted by boiling in 2× Laemmli SDS sample buffer for 3 minutes. Protein samples were separated on pre-cast Bis-Tris polyacrylamide gels (Invitrogen) and stained with Novex colloidal blue staining kit (Invitrogen). Lanes were cut into seven regions according to migration of molecular weight markers and in-gel digestion with trypsin, and mass spectrometry using a 4800 MALDI TOF/TOF Analyser (ABSciex) equipped with a Nd: YAG 355nm laser was performed by St. Andrews University Mass Spectrometry and Proteomics Facility. Mass spectrometry data was analysed using the Mascot search engine (Matrix Science) to interrogate the NCBInr database using tolerances of ± 0.2 Da for peptide and fragment masses, allowing for one missed trypsin cleavage, fixed cysteine carbamidomethylation and variable methionine oxidation.

#### Size Exclusion Chromatography

Superdex 200 10/300 GL (GE Healthcare Life Sciences) was calibrated with molecular weight markers for gel filtration (Sigma-Aldrich) in BC200 buffer (25 mM HEPES pH 7.3, 200 mM NaCl, 1 mM MgCl_2_, 0.5 mM EGTA, 0.1 mM EDTA, 10% glycerol, 1 mM DTT, and 0.2 mM PMSF). 2 mg cytoplasmic extract from ESCs grown in LIF+serum were diluted in 500 µl buffer A/B (15 mM HEPES pH7.6, 115 mM KCl, 3 mM MgCl_2_, 0.1 mM EDTA, 1 mM DTT, 0.2 mM PMSF, Complete protease inhibitors (Roche)) containing 20 µg RNase Inhibitor (Promega), centrifuged (12,000*g*, 10 minutes at 4°C), then loaded on the column. The column was run isocratically in BC200 buffer for 1.4 column volumes and 0.5 ml fractions were collected. Fractions were precipitated with trichloroacetic acid and resuspended in Laemmli SDS sample buffer.

#### Co-Immunoprecipitation

HEK293T cells were transfected with plasmids (pCAG-*Tex19.1*-YFP, pCAG-*TEX19*-YFP, pEGFP3N1-*Ubr2*, pCMV5-hORF1-T7, pCMV5-mORF1-T7, pCMV5-mORF1-mCherry, pCMV5-hORF1^RA^-T7, pCMV5-mORF1^RA^-T7, Supplementary Table 3) using Lipofectamine 2000 (Invitrogen) according to the manufacturer’s instructions and incubated for 24 hours before harvesting. GFP-Trap agarose beads (Chromotek) were used to immunoprecipitate YFP-or GFP-tagged proteins following manufacturer’s instructions. RFP-Trap agarose beads (Chromotek) was similarly used to immunoprecipitate mCherry-tagged proteins, with the addition of a pre-clearing step using binding control agarose beads (Chromotek). The ORF1^RA^ mutants contain two mutations in the RNA binding domain of L1-ORF1p (R260A and R261A in hL1-ORF1p, R297A and R298A in mL1-ORF1p) that reduce the ability of L1-ORF1p to bind RNA and form a RNP (Kulpa and Moran 2005; Martin et al. 2005). These mutations abolish the ability of engineered L1 reporters to retrotranspose (Supplementary Figure S2F) (Moran et al. 1996).

For anti-FLAG immunoprecipitation, cell pellets were lysed for 20 minutes on ice in lysis buffer (10 mM Tris pH 7.5, 150 mM NaCl, 0.5 mM EDTA, 0.5% NP-40, 1 mM PMSF, Complete Protease Inhibitors (Roche)), and insoluble material removed by centrifugation at 12,000*g* for 10 minutes at 4°C. Supernatants were diluted 1:4 in lysis buffer without NP-40, then combined with washed anti-FLAG M2 affinity gel (Sigma), and rotated at 4°C for 1 hour. The anti-FLAG gel was washed three times in lysis buffer without NP-40, then protein eluted in 2× Laemmli SDS sample buffer for Western blot analysis.

#### *In Vivo* Ubiquitylation Assays

HEK293T cells were cotransfected with equal amounts of the indicated plasmids (pCMV-*TEX19*, pCMV-His_6_-myc-ubiquitin (Ward et al. 1995), and pCMV5-hORF1-T7, Supplementary Table 3) using Lipofectamine 2000 (Invitrogen). Cells were harvested 72 hours after transfection and lysed in 6 M guanidinium–HCl, 0.1 M Na_2_HPO_4_, 0.1 M NaH_2_PO_4_, 0.01 M Tris–HCl pH 8.0, 5 mM imidazole and 10 mM β-mercaptoethanol. Following sonication, samples were rotated with washed Ni-NTA agarose (Qiagen) at room temperature for 4 hours. The agarose beads were washed as described (Rodriguez et al. 1999) and ubiquitylated proteins eluted with 200 mM imidazole, 0.15 M Tris–HCl pH 6.7, 30% glycerol, 0.72 M β-mercaptoethanol and 5% SDS then analysed by Western blotting.

#### TUBE2 Pull-Downs

E14 cells were lysed (50 mM Tris pH 7.5, 0.15 M NaCl, 1 mM EDTA, 1% NP-40, 10% glycerol, 5 mM N-ethylmaleimide, Complete Protease Inhibitors (Roche)) on ice for 20 minutes. Cell lysates were centrifuged at 12,000*g* for 10 minutes at 4°C and soluble supernatant incubated at 4°Covernight with TUBE2 or control agarose (LifeSensors) prepared according to manufacturer’s instructions. Agarose beads were washed three times in 50 mM Tris pH 7.4, 150 mM NaCl, 0.1% Tween and protein eluted with 2× Laemmli SDS sample buffer.

#### Strep Pull-Down From Bacterial Lysates

For the Strep pull-down assays, tagged hL1-ORF1p and human TEX19 were either co-expressed or separately expressed overnight at 20 °C in *Escherichia coli* BL21 (DE3) Star cells. The cells were lysed in a binding buffer (50 mM HEPES pH 7.0, 200 mM NaCl, 2 mM DTT) containing DNase I, lysozyme and protease inhibitors. For proteins expressed separately, 200 µl of the Strep-tagged binding partner (ORF1p or GB1) were incubated with 50 µl Strep-Tactin Sepharose beads (IBA) in a total volume of 1 ml of binding buffer for 45 minutes at 4 °C. After centrifugation (~1500 g) and two washes with 700 µl of binding buffer, 1 ml of TEX19 lysate was added to the beads, followed by an additional incubation for 45 minutes at 4 °C. For co-expressed proteins, 1 ml of the lysate was added to 50 µl Strep-Tactin Sepharose beads (IBA) and incubated for 45 minutes at 4 °C. In the end, the beads were washed five times with 700 µl of binding buffer. The bound proteins were eluted with 100 µl of the binding buffer supplemented with 2.5 mM biotin. The eluted proteins were then precipitated by trichloroacetic acid, resuspended in Laemmli SDS sample buffer and analyzed by SDS-PAGE.

#### Luciferase Assays

Luciferase activity was measured 24 hours post-transfection using the Dual-Luciferase Reporter Assay system (Promega) following manufacturer’s instructions.

#### Retrotransposition Assays

We used three different L1 retrotransposition assays in HEK293T, U2OS, HeLa and mouse ESCs. In all retrotransposition assays, we also included a transfection efficiency control to calculate rates of engineered retrotransposition as described (Garcia-Perez et al. 2010; Kopera et al. 2016). Where indicated, engineered L1 retrotransposons were co-transfected with a second expression plasmid for *TEX19* orthologs or controls. For *mneoI* and *mblastI*-based assays, we included a plasmid containing a neomycin or blasticidin resistance expression cassette respectively, to control for cytotoxicity (Kopera et al. 2016; Richardson et al. 2014b) when over-expressing *TEX19* orthologs.

Retrotransposition assays with *mneoI* or *mblastI* tagged L1 constructs in cultured HeLa and U2OS cells were performed as described (Kopera et al. 2016; Morrish et al. 2002; Richardson et al. 2014b; Wei et al. 2000). HeLa cells were transfected with Fugene6 (Promega) using 1 μg plasmid DNA per 35 mm diameter well and OptiMEM (Invitrogen) according to the manufacturer instructions. 400 μg/ml G418 selection for 12 days was initiated 72 hours post-transfection for *mneoI* constructs, or 10 μg/ml blasticidin S selection was initiated 120 hours post-transfection for 7 days for *mblastI* constructs. Drug-resistant foci were then fixed (2% formaldehyde, 0.2% glutaraldehyde in PBS) and stained (0.1% crystal violet). Retrotransposition assays with *mneoI* or *mblastI* tagged L1 constructs in mouse ESCs were conducted by plating 4×10^5^ cells per 35 mm diameter well onto gelatin-coated tissue culture plates and transfecting 18 hours later with Lipofectamine 2000 (Invitrogen) using 1 μg plasmid DNA per well and OptiMEM (Invitrogen) according to the manufacturer instructions. Media was replaced after 8 hours and transfected mouse ESCs passaged into a gelatin-coated 100mm tissue culture plate 24 hours later. 200 μg/ml G418 or 8 μg/ml blasticidin S selection for 12 days was initiated after an additional 24 hours, and drug-resistant foci fixed, stained and counted as described for HeLa cells.

Retrotransposition assays with mEGFPI tagged L1 constructs in cultured HEK293T cells were performed as described (Goodier et al. 2013; Wei et al. 2000). 2×10^5^ HEK293T cells were plated in a 35 mm diameter well, then transfected with Lipofectamine 2000 (Invitrogen) and 1μg plasmid DNA per well using OptiMEM (Invitrogen) following the manufacturer instructions 20 hours later. 24 hours later, fresh media containing 2 μg/ml puromycin (Sigma) was added daily for 7 days. Cells were collected by trypsinization and the percentage of EGFP-expressing cells determined using a FACSCanto II flow cytometer (BD Biosciences). Transfection with mutant L1 plasmid (99-gfp-JM111 or 99-gp-L1SMmut2) allowed a threshold to be established for background fluorescence.

#### Confocal Microscopy

1×10^5^ U2OS cells were plated in 35 mm diameter wells, then 20 hours later transfected with Fugene6 (Promega) and 1μg plasmid DNA per well using OptiMEM (Invitrogen) following the manufacturer instructions. Media was replaced 20 hours after transfection and cells allowed to grow for a total of 36 hours. Next, the transfected cells were trypsinised and 25-50% plated on a 15 mm diameter sterile circular polysterene coverslip in a 35 mm diameter well. 12 hours later, cells were fixed with 4% paraformaldehyde at room temperature for 30 minutes, permeabilised with PBS containing 0.1% (v/v) Triton X-100, then incubated with blocking solution (1% normal goat serum, in PBS) for 30 minutes. After two washes in PBS containing 0.1% goat serum and 0.05% Triton X-100, coverslips were incubated with 1:600 rabbit anti-T7 primary antibody (Abcam) diluted in PBS at 4°C overnight in a humidified chamber. Coverslips were then washed three times and incubated with 1:1000 Alexa Fluor-conjugated anti-rabbit secondary antibodies for 30 minutes at room temperature. Coverslips were then washed twice and mounted with SlowFade Gold antifade with DAPI (ThermoFisher) and sealed with nail polish. Slides were imaged using a Zeiss LSM-710 confocal microscope (Leica), an Axio Imager A1 Microscope (Zeiss) and captured images analyzed with ZEN lite software (Zeiss).

## 7 Supplementary Information

### Supplementary Tables

**Table 1.**
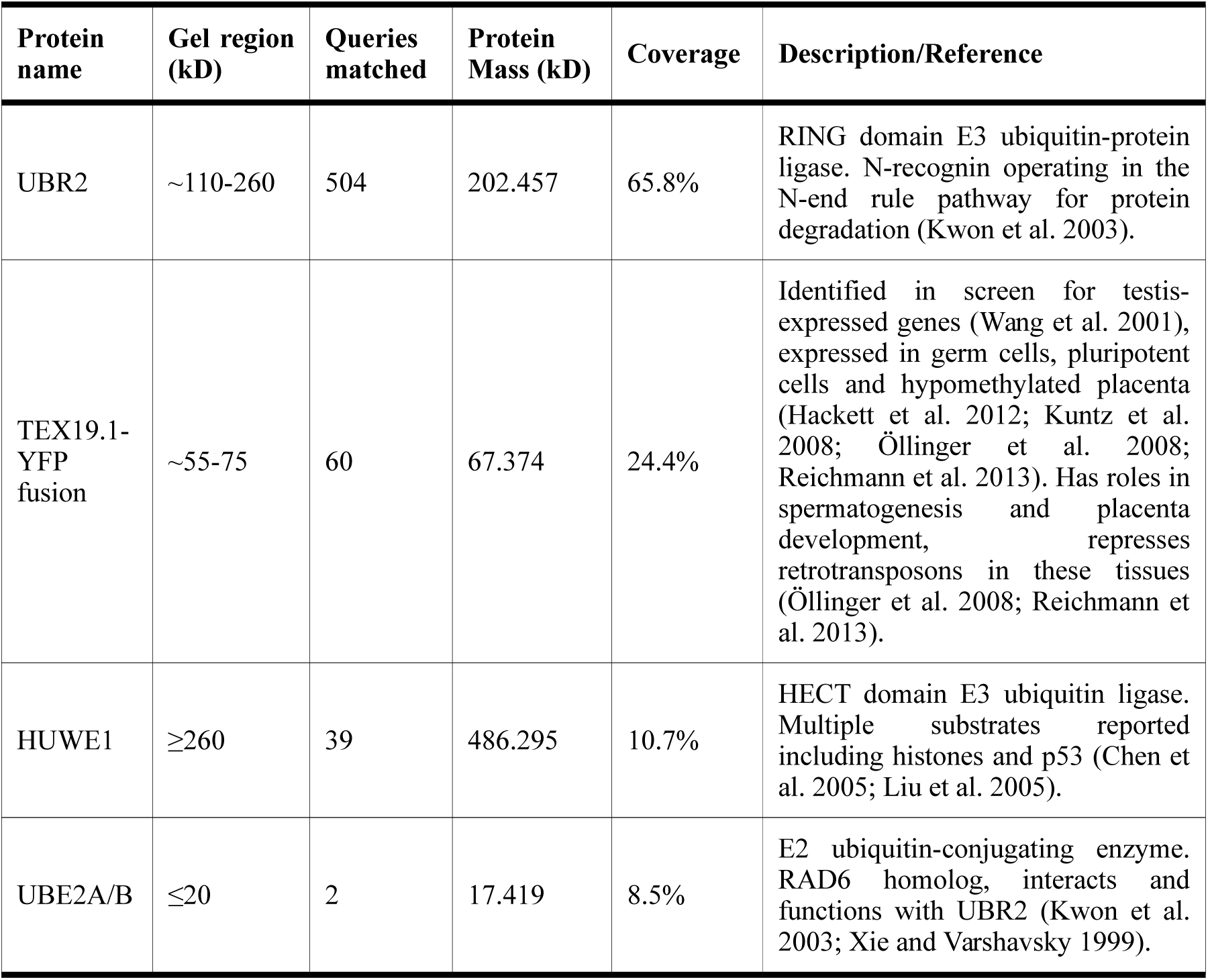
**Proteins Identified In TEX19.1-YFP Immunoprecipitates.** Proteins identified by mass spectrometry in TEX19.1-YFP immunoprecipitates from mouse ES cell cytoplasmic lysates, but not in YFP controls. Only interactors verified by Western blotting (Figure 2 are listed. Queries matched indicates the number of MS/MS spectra that were matched to each protein, coverage indicates the percentage of target protein matched by MS/MS spectra.

**Table 2.**
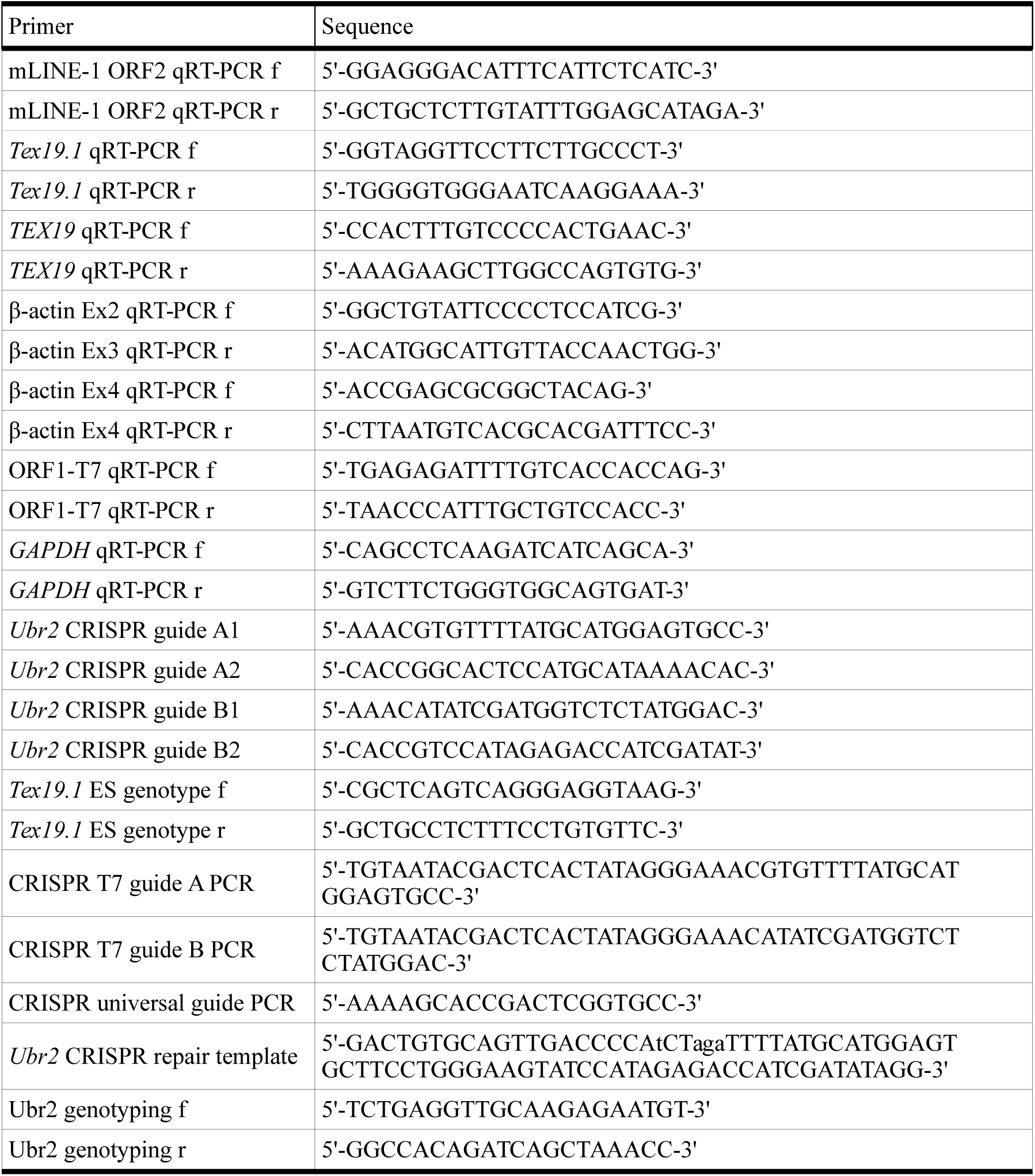
**Oligonucleotides Used In This Study.**Lower case nucleotides in the repair template sequence indicate mutations relative to wild-type sequence.

**Supplementary Table 3.** Plasmids Used In This Study.

| Plasmid | Description |
| --- | --- |
| 99-gfp-LRE3 | Described previously (Coufal et al. 2009), contains a full-length retrotransposition-competent human L1 element (LRE3) (Brouha et al. 2002) tagged with the *mEGFP* retrotransposition indicator cassette (Ostertag et al. 2000) in pCEP4 (Invitrogen) lacking a CMV promoter and containing a puromycin rather than a hygromycin resistance cassette (Garcia-Perez et al. 2010). |
| 99-gfp-JM111 | Described previously (Coufal et al. 2009). A derivative of 99-gfp-LRE3 where the cloned human L1, L1RP (Kimberland et al. 1999), contains two engineered missense mutations in hL1-ORF1p (R261A, R262A). |
| 99-gfp-TGF21 | Described previously (Garcia-Perez et al. 2010), contains a full-length retrotransposition-competent mouse L1GF element (TGF21) (Goodier et al. 2001) tagged with the *mEGFP* retrotransposition indicator cassette (Ostertag et al. 2000) cloned in pCEP4 (Invitrogen) lacking a CMV promoter and containing a puromycin rather than a hygromycin resistance cassette (Garcia-Perez et al. 2010). |
| 99-gfp-L1SM | Described previously (Garcia-Perez et al. 2010), contains a full-length retrotransposition-competent codon-optimised mouse L1 T<sub>F</sub> element (L1spa codon-optimised) (Han and Boeke 2004) tagged with the *mEGFP* retrotransposition indicator cassette (Ostertag et al. 2000) cloned in pCEP4 (Invitrogen) lacking a CMV promoter and containing a puromycin rather than a hygromycin resistance cassette (Garcia-Perez et al. 2010). |
| 99-gfp-L1SMmut2 | Described previously (Garcia-Perez et al. 2010), a derivative of 99-gfp-L1SM where the cloned mouse L1 contains two missense mutations in the endonuclease and reverse transcriptase domains of ORF2p (D212G and D709Y respectively) (Han and Boeke 2004). |
| pCEPL1SM | Described previously (Han and Boeke 2004), contains a full-length retrotransposition-competent codon-optimised mouse L1 T<sub>F</sub> element (L1spa codon-optimised) (Han and Boeke 2004) tagged with the *mneoI* retrotransposition indicator cassette (Freeman et al. 1994) and cloned in pCEP4 (Invitrogen). |
| pCEPL1SM-T7 | A derivative of pCEPL1SM where a T7-epitope tag has been cloned in the C-terminus of ORF1p. Also, the L1 element lacks the 5' UTR from L1spa. |
| pCEPL1SM-T7-ORF1<sup>RA</sup> | A derivative of pCEPL1SM-T7 that contains two missense mutations in the RNA binding domain of ORF1p (R297A, R298A). These mutations abolish the ability of the codon-optimised L1 T<sub>F</sub> element to mobilise. |
| pAD2TE1 | Described previously (Doucet et al. 2010), contains a full-length retrotransposition-competent human L1 (L1.3) (Sassaman et al. 1997) tagged with the *mneoI* retrotransposition indicator cassette (Freeman et al. 1994) and cloned pCEP4 (Invitrogen). A T7 and a TAP epitope tag have been cloned in the C-terminus of ORF1p and ORF2p respectively. |
| pCEPL1SMN21A | Described previously (Alisch et al. 2006), a derivative of pCEPL1SM that contains a missense mutation in the endonuclease domain of ORF2p (N21A). |
| Zfl2-2mneoI | Described previously (Sugano et al. 2006), contains a full-length retrotransposition-competent zebrafish LINE-2 element tagged with the *mneoI* retrotransposition indicator cassette (Freeman et al. 1994) and cloned in pCEP4 (Invitrogen). |
| pU6ineo | Described previously (Richardson et al. 2014b), contains the neomycin phosphotransferase expression cassette from pEGFP-N1 (Clontech) cloned in a pBluescript II KS (pBSKS, Stratagene). |
| JJ101/L1.3 | Described previously (Beck et al. 2010), contains a full-length retrotransposition-competent human L1 (L1.3) (Sassaman et al. 1997) tagged with the *mblastI* retrotransposition indicator cassette (Goodier et al. 2007) and cloned in pCEP4 (Invitrogen). |
| JJL1SM | Contains a full-length retrotransposition-competent mouse L1 T<sub>F</sub> element (L1spa codon-optimised) (Han and Boeke 2004) tagged with the *mblastI* retrotransposition indicator cassette (Goodier et al. 2007) and cloned in pCEP4 (Invitrogen). |
| pCMV5-hORF1-T7 | Described previously (Kulpa and Moran 2005), contains the coding sequence of L1.3 hL1-ORF1p (Sassaman et al. 1997) tagged with a C-terminal T7 epitope in vector pCMV5. |
| pCMV5-hORF1-RA-T7 | A derivative of pCMV5-hORF1-T7 where the coding sequence of ORF1p contains two missense mutation in the RNA binding domain (R261A, R262A). |
| pCMV5-mORF1-T7 | Contains the coding sequence of ORF1p from the codon-optimised mouse L1 T<sub>F</sub> element (Han and Boeke 2004) tagged with a C-terminal T7 epitope in vector pCMV5. |
| pCMV5-mORF1- RA-T7 | A derivative of pCMV5-mORF1-T7 where the coding sequence of ORF1p contains two missense mutation in the RNA binding domain (R297A, R298A). |
| pCMV5-hORF1-mCherry | Contains the coding sequence of L1.3 hL1-ORF1p (Sassaman et al. 1997) with a C-terminal mCherry tag (Shaner et al. 2005) in vector pCMV5. |
| pCMV5-mORF1-mCherry | Contains the coding sequence of codon-optimised mouse L1 T<sub>F</sub> element (Han and Boeke 2004) with a C-terminal mCherry tag (Shaner et al. 2005) in vector pCMV5. |
| pK-APOBEC3A | Described previously (Bogerd et al. 2006a), expression vector for APOBEC3A. |
| pK-β-ARRESTIN | Described previously (Bogerd et al. 2006a), expression vector for β-ARRESTIN. |
| pGL3SV40LUC | Obtained from Promega, luciferase under control of a SV40 promoter. |
| pTKLUC | Obtained from Promega, luciferase under control of a HSV-TK promoter. |
| pL1.3FF | Described previously (Heras et al. 2013), contains 5'UTR from a human L1.3 L1 element (Sassaman et al. 1997) cloned in pGL3-Basic luciferase reporter vector (Promega). |
| pL1spaFF | Described previously (Heras et al. 2013), contains 5'UTR from a mouse L1 T<sub>F</sub> element (L1spa) cloned into pGL3-Basic (Promega) |
| pCEP-EGFP | Described previously (Alisch et al. 2006), has the enhanced green fluorescent protein (EGFP) coding sequence in pCEP4 (Invitrogen). |
| pcDNA6.1/mycA | Obtained from Invitrogen. contains a blastidicin resistance cassette. |
| pBSKS | Obtained from Stratagene, pBluescript II KS phagemid cloning vector. |
| pCAG-YFP | The CMV promoter in pEYFP-N1 (Clontech) was replaced with the CAG promoter from pCAGGS (Niwa et al. 1991). |
| pCAG-TEX19.1-YFP | The mouse TEX19.1 open reading frame was inserted into the pCAG-YFP plasmid to tag TEX19.1 on the C-terminus with EYFP. |
| pCAG-TEX19-YFP | The human TEX19 open reading frame was inserted into the pCAG-YFP plasmid to tag TEX19 on the C-terminus with EYFP. |
| pCMV-TEX19 | Human TEX19 under control of a CMV promoter in pCMV-SPORT6 |
| p105 | Obtained from IBA Life Sciences, pEXPR-IBA105 eukaryotic expression vector with CMV promoter and Strep-Tactin (Strep) tag. |
| p105-TEX19.1 | Mouse *Tex19.1* coding sequence cloned into p105 to generate a Strep tag at the N-terminus. |
| p105-TEX19 | Human *TEX19* coding sequence cloned into p105 to generate a Strep tag at the N-terminus. |
| p3xFLAG-CMV-Ubr2 | Mouse *Ubr2* coding sequence cloned in to p3XFLAG-CMV-10 (Sigma) expression vector that generate a 3xFLAG epitope tag at the N-terminus of UBR2. |
| pEGFP3N1-Ubr2 | Mouse *Ubr2* coding sequence cloned into pEGFP-N1 (Clontech) expression vector to generate a GFP tag at the C-terminus of UBR2. |
| pCMV-His<sub>6</sub>-myc-ubiquitin | Described previously (Ward et al. 1995), vector expressing ubiquitin tagged with hexahistidine and myc epitopes. |
| pRF | Described previously (Li et al. 2006), dicistronic reporter construct containing Renilla and firefly luciferase coding sequences. |
| pRFA | Described previously (Li et al. 2006), 400 bp upstream of mL1-ORF1p (L1spa 5' UTR) inserted between luciferase coding sequences in pRF. |
| pRFD | Described previously (Li et al. 2006), 200 bp upstream of mL1-ORF2p (L1spa intergenic region) inserted between luciferase coding sequences in pRF. |
| pRF3 | Described previously (Li et al. 2006), 312 bp from the 3'UTR of mouse L1 (L1spa 3'UTR) inserted between luciferase coding sequences in pRF. |
| pnEA-pS-hL1ORF1p (AII) | hL1-ORF1p coding sequence from pJM101 (Moran et al. 1996) inserted into pnEA-pS (Diebold et al. 2011) for bacterial expression and to generate a cleavable Strep tag at the N-terminus (engineered point mutations: M121A, M125I, M128I) (Khazina et al. 2011). |
| pnEA-pS-B | GB1 coding sequence (Cheng and Patel 2004) inserted into pnEA-pS (Diebold et al. 2011) for bacterial expression and to generate a Strep tag at the N-terminus. |
| pnYC-pM-hTex19-pBH | Human *TEX19* coding sequence from pCMV-TEX19 inserted into pnYC-pM (Diebold et al. 2011) for bacterial expression and to generate a cleavable MBP tag at the N-terminus. A cleavable C-terminal GB1 tag (Cheng and Patel 2004) was also introduced, followed by a hexahistidine tag. |

**Supplementary Table 4.** Antibodies Used For Western Blots.

| **Antibody** | **Source** | **Dilution** |
| --- | --- | --- |
| Rabbit anti-TEX19.1 | Ian Adams (Öllinger et al. 2008) | 1:100 |
| Rabbit anti-mouse L1 ORF1p | Sandy Martin/Alex Bortvin (Martin and Branciforte 1993; Soper et al. 2008) | 1:2000 |
| Rabbit anti-mouse L1 ORF1p | Oliver Weichenrieder | 1:2000 |
| Mouse anti-β-ACTIN | Sigma (A5441) | 1:5000 |
| Mouse anti-UBR2 | Abcam (ab57407) | 1:1000 |
| Goat anti-UBR2 | Aviva Systems Biology (OAEB00482) | 1:100 |
| Rabbit anti-PABP1 | Niki Gray (Burgess et al. 2011) | 1:10000 |
| Rabbit anti-HUWE1 | Bethyl Laboratories (A300-486A) | 1:500 |
| Rabbit anti-UBE2A | GeneTex (GTX100426) | 1:3000 |
| Mouse anti-GFP/YFP | Roche (118144600001) | 1:2000 |
| Mouse anti-RFP/mCherry | Chromotek (6G6) | 1:5000 |
| Rabbit anti-T7 | Abcam (ab18611) | 1:5000 |
| Rabbit anti-Lamin B1 | Abcam (ab16048) | 1:5000 |
| Mouse anti-c-Myc | Sigma (M4439) | 1:5000 |
| Rabbit anti-TEX19 | Abcam (ab185507) | 1:2500 |
| Rabbit anti-histone H3 | Abcam (ab1791) | 1:20000 |
| Rabbit anti-p53 | Santa Cruz Technologies | 1:300 |

## Supplementary Figure Legends

**Supplementary Figure S1.**
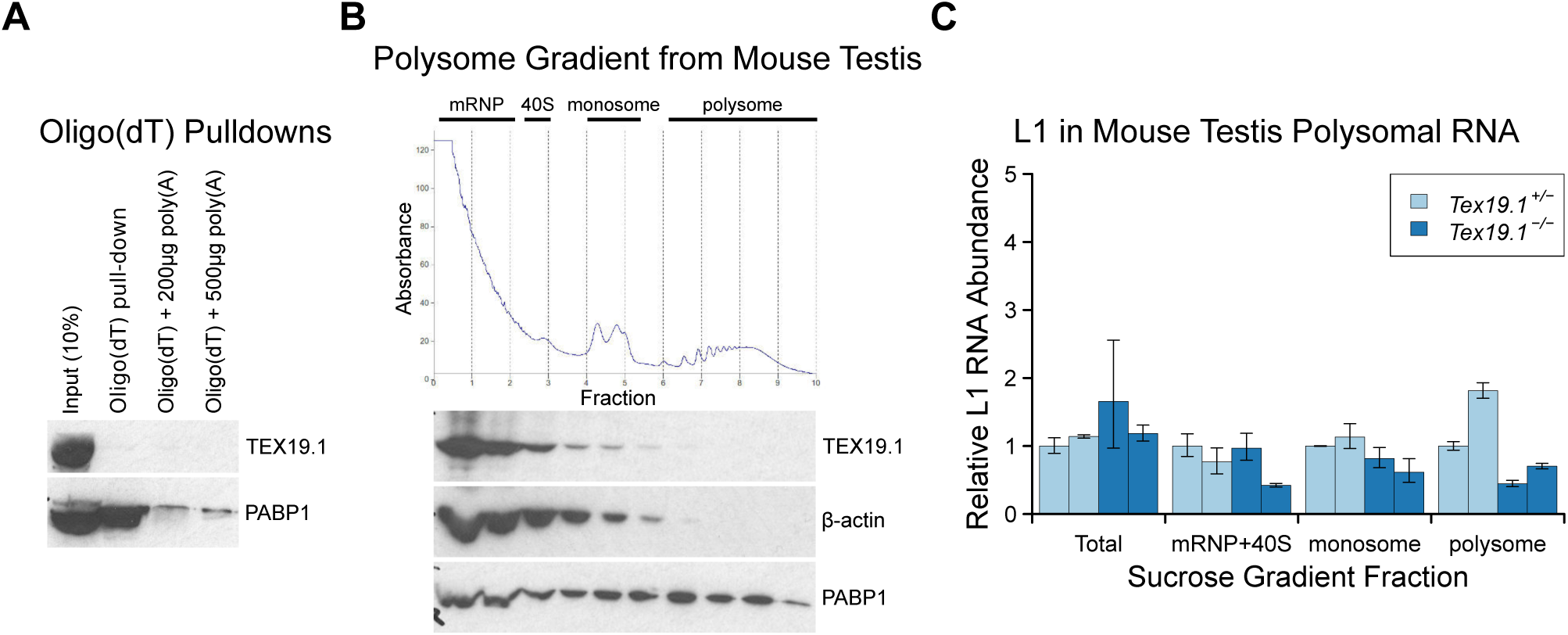
***Tex19.1* Does Not Inhibit L1 Translation. A.** Oligo(dT) pull-downsfrom P16 testes. Oligo(dT) cellulose beads were used to isolate poly(A) RNA from testis lysates, and associated proteins analysed by Western blotting with indicated antibodies. 200 or 500 µg poly(A) RNA was added as a competitor. The poly(A) RNA binding protein PABP1 was used as a positive control. TEX19.1 is not detectably associated with poly(A) RNA in testes. **B.** Sucrose density gradient enrichment of translation intermediates from P18 testes. The protein content of the fractions was monitored by reading absorbance at 254 nm, and peaks corresponding to messenger ribonucleoproteins (mRNPs), 40S ribosomal subunits, monosomes and polysomes are indicated. Western blots for TEX19.1, β-actin and PABP1 are shown for each fraction. TEX19.1 is not detectably associated with actively translating polysomes in testes. **C.** qRT-PCR for L1 mRNA in mRNP+40S, monosome, and polysome fractions in sucrose gradients from *Tex19.1*^*+/−*^ and *Tex19.1*^*−/−*^P18 testes. L1 mRNA abundance was measured relative to β-actin in each fraction, and normalised to one of the heterozygous control animals. A proportion of L1 mRNA associates with polysomes consistent with previous reports (Tanaka et al.2011). Meiotic arrest and increased spermatocyte death between P16 and P22 in *Tex19.1*^*−/−*^ testes (Öllinger et al. 2008) may be generating some differences in testicular cell composition in these P18 samples and causing subtle differences in L1 mRNA distribution between *Tex19.1*^*+/−*^ and *Tex19.1*^*−/−*
^ samples. However, there is no statistically significant increase in polysome-associated L1 mRNA in *Tex19.1*^*−/−*^ P18 testes. Error bars indicate SEM for technical replicates.

**Supplementary Figure S2.**
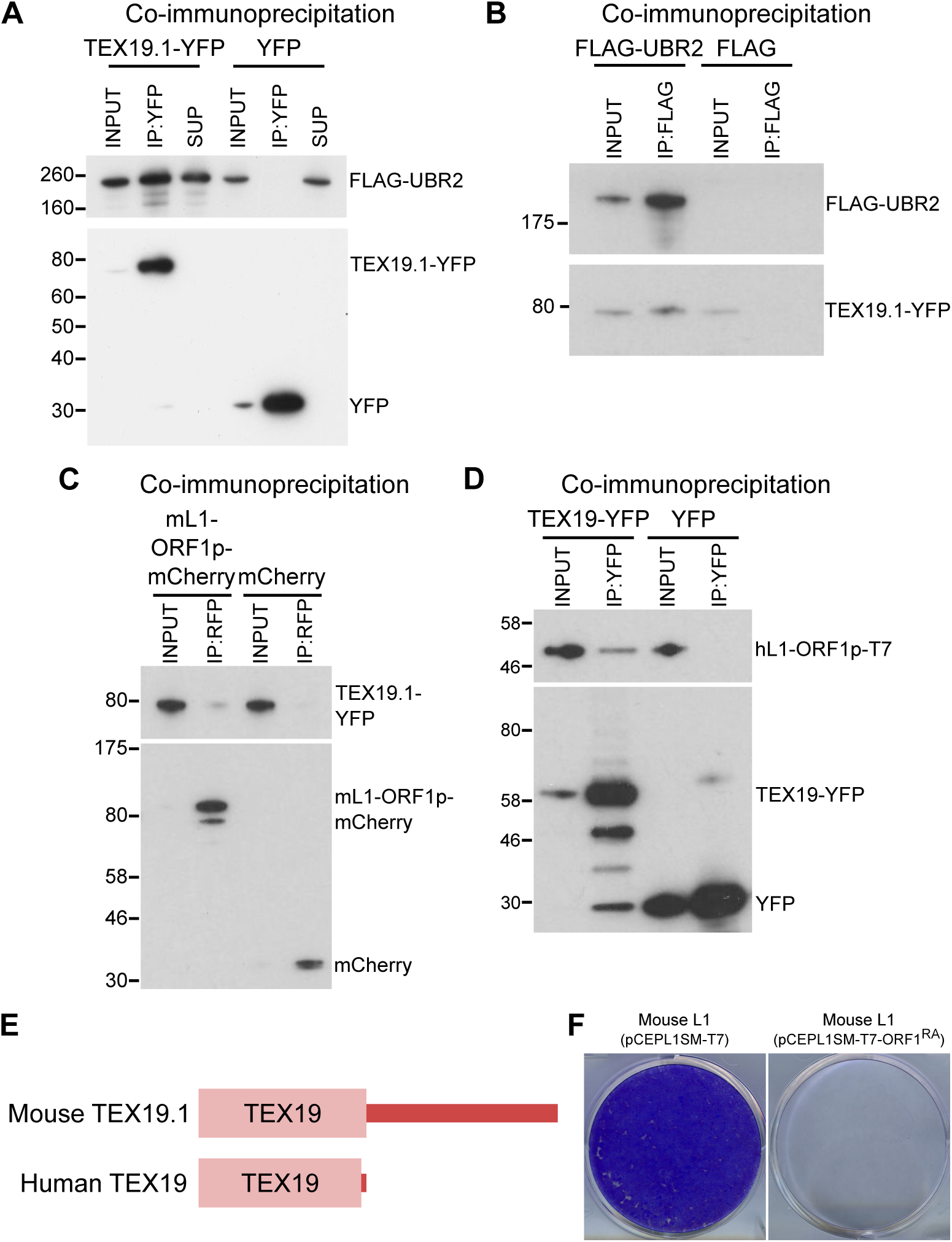
**TEX19 Orthologs Interact With UBR2 And L1-ORF1p. A.** Co-immunoprecipitations (co-IPs) from stable HEK293 cell lines expressing either TEX19.1-YFP or YFP alone transiently transfected with FLAG-UBR2. Anti-YFP immunoprecipitates (IPs), inputs, and supernatants (SUP) were Western blotted with anti-FLAG and anti-YFP antibodies. **B.** Reciprocal co-IP for panel A. HEK293T cells were transiently transfected with TEX19.1-YFP and either FLAG-UBR2 or FLAG alone, and anti-FLAG IPs and their inputs were Western blotted with anti-FLAG and anti-YFP antibodies. Positions of FLAG-UBR2, TEX19.1-YFP, YFP alone and pre-stained molecular weight markers in kD are indicated. **C.** Co-immunoprecipitation from HEK293T cells co-transfected with TEX19.1-YFP and mCherry-tagged mL1-ORF1p expression constructs and IPd for mCherry. YFP or mCherry alone were used as negative controls. Anti-mCherry IP inputs and IPs were Western blotted with anti-mCherry or anti-YFP antibodies. Positions of pre-stained molecular weight markers in kD are indicated. **D.** Co-IPs from HEK293T cells co-transfected with epitope-tagged hL1-ORF1p and human TEX19-YFP expression constructs. YFP was used as a negative control. IP inputs and IPs were Western blotted with anti-T7 or anti-YFP antibodies. **E.** Diagram showing the domain structure of mouse and human TEX19 orthologs. A conserved TEX19 domain is present at the N-terminus of both proteins, but the C-terminal region of mouse TEX19.1 is not conserved in the truncated human TEX19 protein. **F.** Mouse L1-ORF1^RA^p mutants used to test for RNA-independent interactions have impaired mobilisation. Plates of G418-resistant colonies from L1 retrotransposition assays in HeLa cells. Assays for mouse L1 (pCEPL1SM-T7) and mouse L1 carrying the R297A and R298A mutations in the RNA binding domain of ORF1p that reduce its affinity for RNA (Martin et al. 2005) (pCEPL1SM-T7-ORF1^RA^).

**Supplementary Figure S3.**
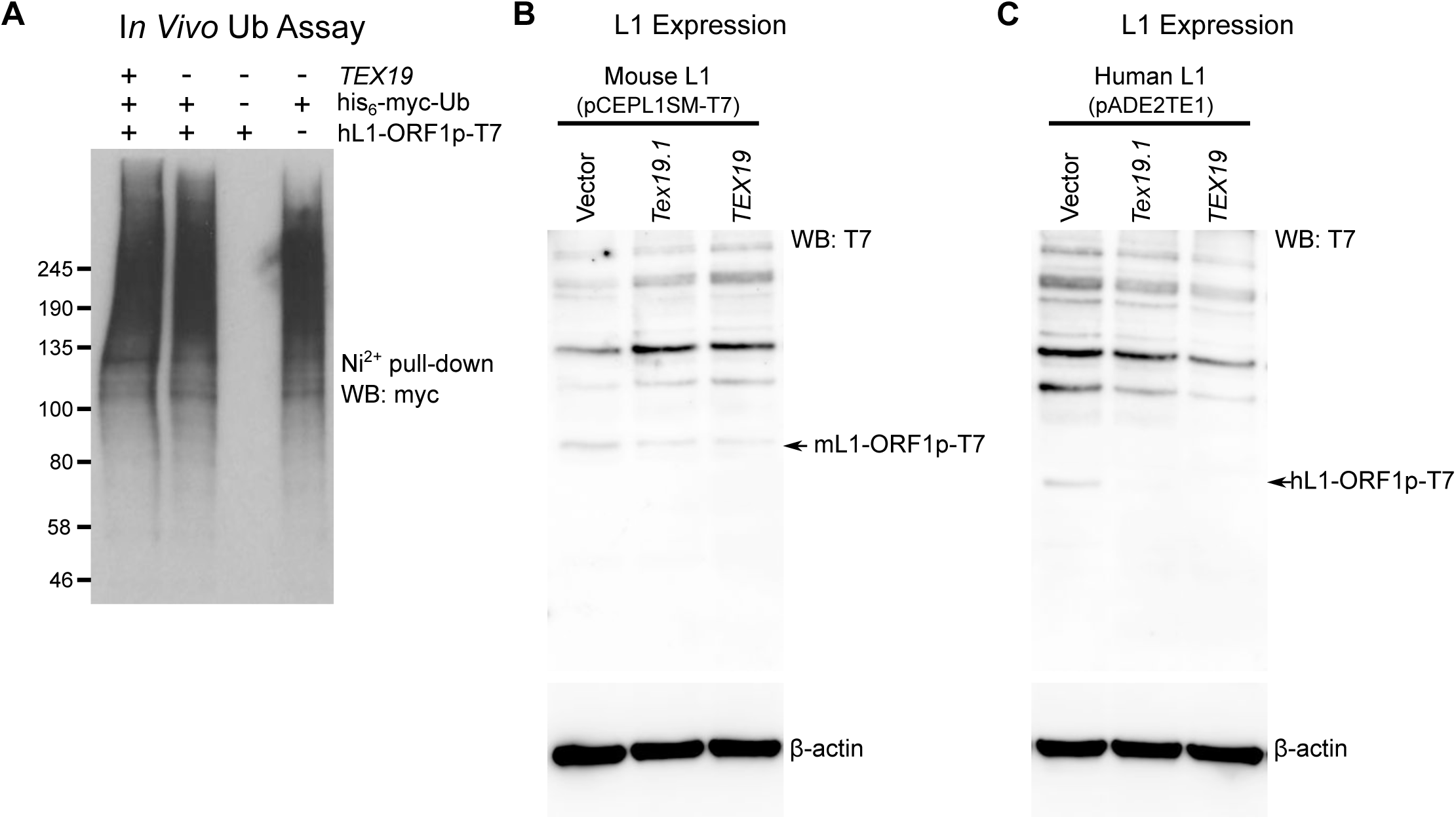
***TEX19* Orthologs Regulate L1-ORF1p Abundance. A.** Control for *in vivo* ubiquitylation assay shown in Figure 3C. Ni^2+^pull-downs were Western blotted (WB) withanti-T7 antibodies to detect the immunoprecipitated his_6_-myc-Ub conjugates. MW markers (kD) are shown beside blots. **B, C.** Hamster XR-1 cells were transiently transfected with a synthetic mouse or human L1 construct containing T7 epitope-tagged ORF1p (panel B: mouse, pCEPL1SM-T7; panel C, pADE2TE1, human), and either empty vector, Strep-Tex19.1 or Strep-TEX19 expression constructs. Cells were Western blotted (WB) for the T7 epitope tag, and for β-actin as a loading control 72 hours post-transfection. Arrows indicate the L1-ORF1p-T7 bands (43 kD for mL1-ORF1p-T7, 40 kD for hL1-ORF1p-T7). ***TEX19*Orthologs Regulate L1-ORF1p Abundance. A.** Control for *in vivo* ubiquitylation assay shown in Figure 3C. Ni^2+^pull-downs were Western blotted (WB) withanti-T7 antibodies to detect the immunoprecipitated his_6_-myc-Ub conjugates. MW markers (kD) are shown beside blots. **B, C.** Hamster XR-1 cells were transiently transfected with a synthetic mouse or human L1 construct containing T7 epitope-tagged ORF1p (panel B: mouse, pCEPL1SM-T7; panel C, pADE2TE1, human), and either empty vector, Strep-Tex19.1 or Strep-TEX19 expression constructs. Cells were Western blotted (WB) for the T7 epitope tag, and for β-actin as a loading control 72 hours post-transfection. Arrows indicate the L1-ORF1p-T7 bands (43 kD for mL1-ORF1p-T7, 40 kD for hL1-ORF1p-T7).

**Supplementary Figure S4.**
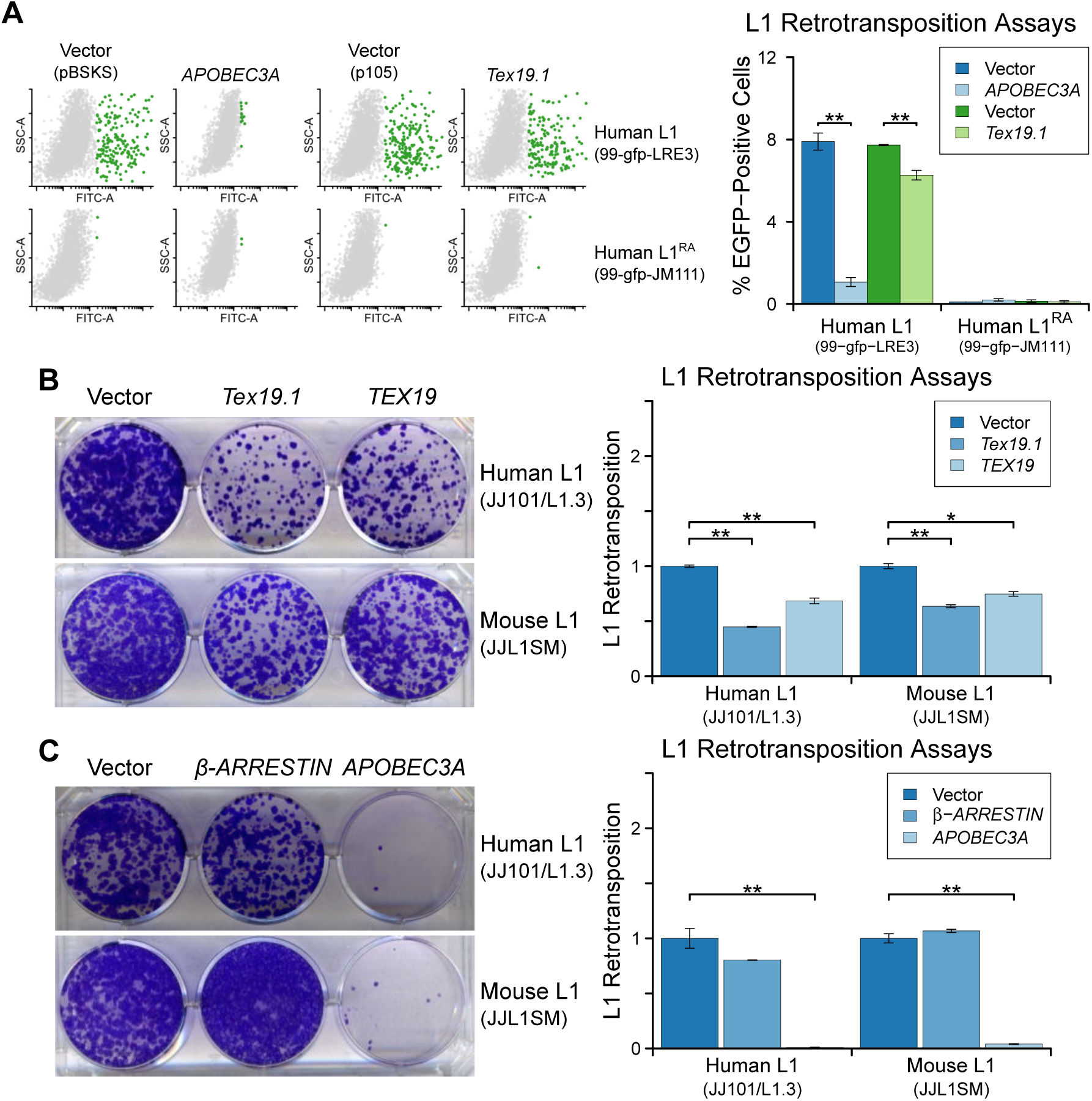
***TEX19* Orthologs Restricting L1 Mobilisation. A.** Flow cytometry profiles of engineered L1 retrotransposition assays in HEK293T cells co-transfected with human L1 constructs (99-gfp-LRE3 and 99-gfp-JM111) containing EGFP retrotransposition reporters, and either Strep-tagged *Tex19.1*, *APOBEC3A* (positive control) or empty vectors (pBSKS for *APOBEC3A*, pIBA105 for *Tex19.1*). EGFP fluorescence is plotted on the x-axis and side scatter onthe y-axis of the flow cytometry profiles, and cells classed as EGFP-positive are shown in green. 99-gfp-JM111 carries the ORF1^RA^ mutations and is severely impaired for retrotransposition (Han and Boeke 2004; Moran et al 1996). *;*; p<0.01 (*t*-test). **B, C.** Blasticidin-resistant colonies from L1 retrotransposition assays in U2OS cells. Human (JJ101/L1.3) and mouse (JJL1SM) engineered L1 constructs containing blasticidin-resistance retrotransposition reporters were co-transfected with Strep-tagged mouse *Tex19.1*, Strep-tagged human *TEX19*, or empty vector (B), or with β*-ARRESTIN* (negative control), *APOBEC3A* (positive control) or empty vector (C). Quantification ofL1 retrotransposition normalised for transfection efficiency is shown. * p<0.05; *;*; p<0.01 (*t*-test).

**Supplementary Figure S5.**
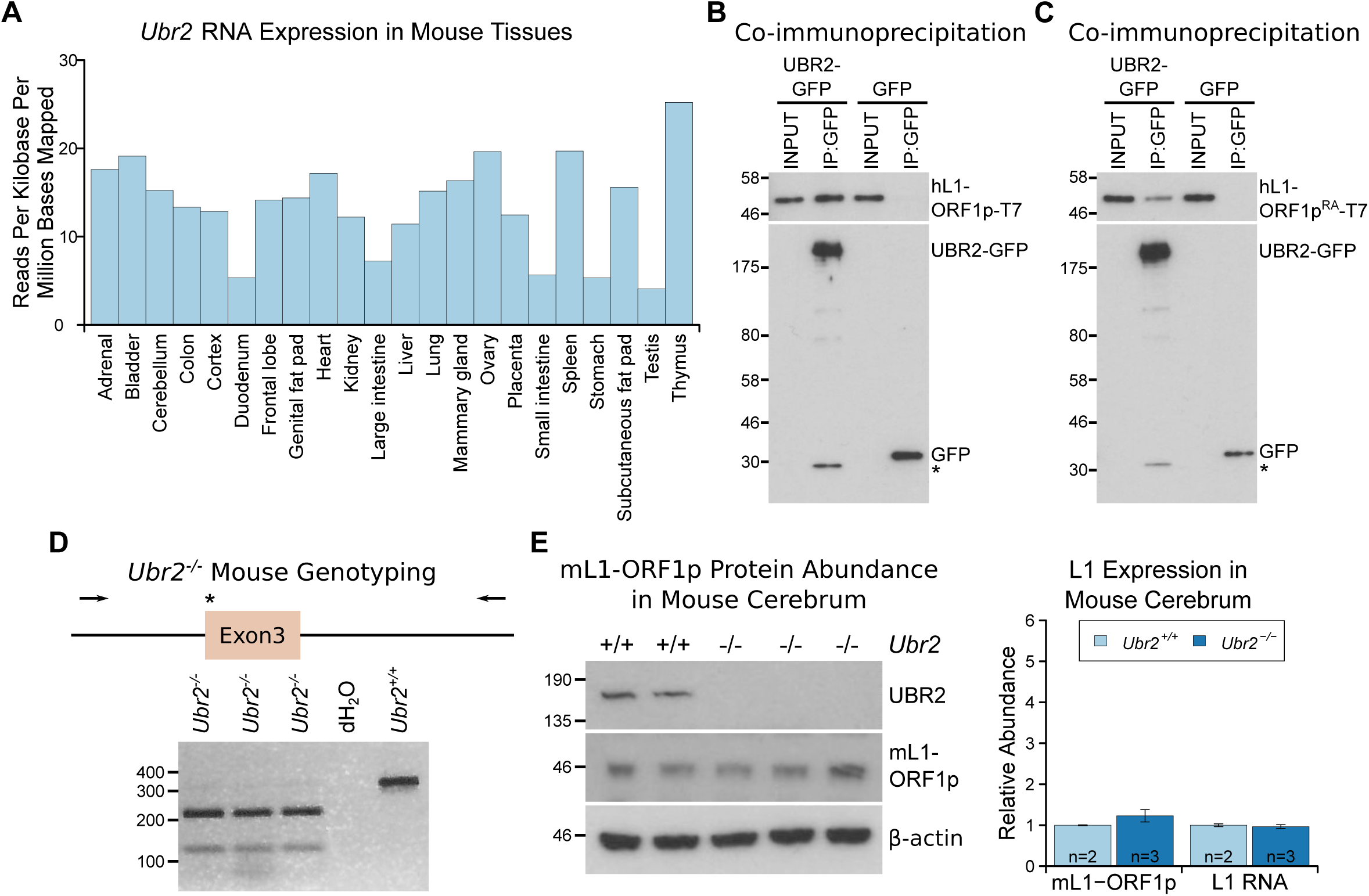
**The Ubiquitously-Expressed E3 Ubiquitin Ligase UBR2 Physically Interacts With L1-ORF1p But Does Not Regulate Its Abundance In The Cerebrum. A**. *Ubr2* transcript abundance in multiple adult tissues was determined from ENCODE RNA sequencing data GSE36025 (Lin et al 2014) by calculating the total number of reads mapped to the *Ubr2* locus per million reads mapped in the dataset, and normalising this to the length of the *Ubr2* locus. B, C. Co-immunoprecipitations (co-IPs) from HEK293T cells co-transfected with T7 epitope-tagged hL1-ORF1p and mouse UBR2-GFP expression constructs. GFP alone was used as a negative control. D. Genotyping of *Ubr2*^*−/−*^ mice. An *Xba*I restriction site and premature stop codon (asterisk) are introduced into exon 3 of *Ubr2* by CRISPR/Cas9, and mice genotyped by amplifying a region encompassing exon 3 (primers indicated by arrows) and digesting the PCR product with *Xba*I. Three *Ubr2*^*−/−*^ mice, and *Ubr2*^*+/+*^ and distilled water (dH_2_O) controls are shown. E. Western blots showing endogenous UBR2 and mL1-ORF1p expression in *Ubr*^*+/+*^ and *Ubr2*^*−/−*^ mouse cerebrum. β-actin was used as a loading control. Positions of epitope-tagged proteins and pre-stained molecular weight markers in kD are indicated. Quantification of endogenous mL1-ORF1p abundance and L1 RNA abundance relative to β-actin in *Ubr2*^*+/+*^ and *Ubr2*^*−/−*^ mouse cerebrum is also shown. Relative abundance was normalised to the mean of the *Ubr2*^*+/+*^ control mice. Error bars indicate SEM, MW markers (kD for protein, bp for DNA) are shown beside blots and gels.

**Supplementary Figure S6.**
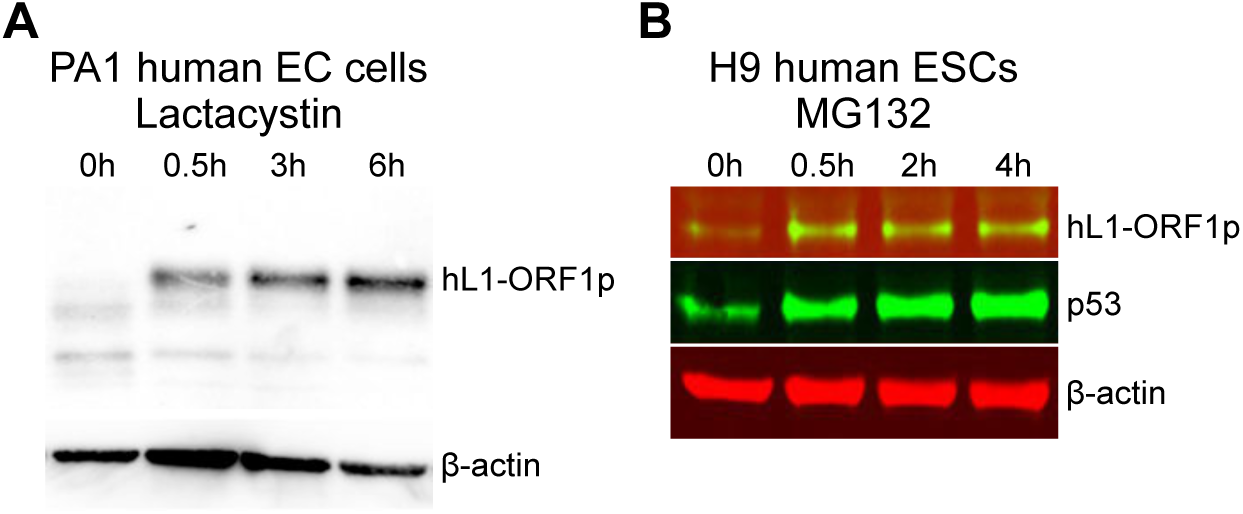
**hL1-ORF1p Abundance In Human Embryonal Carcinoma Cells And Human ESCs Increases In Response To Inhibition Of The Proteasome. A**. Western blot showing abundance of endogenous hL1-ORF1p in PA-1 human embryonal carcinoma (EC) cells after addition of 25 µM lactacystin to inhibit the proteasome. β-actin is shown as a loading control. B. Western blot showing abundance of endogenous hL1-ORF1p in H9 human ESCs after addition of 25 µM MG132 to inhibit the proteasome. p53 is a positive control and accumulates upon MG132 treatment, β-actin is shown as a loading control.

**Supplementary Figure S7.**
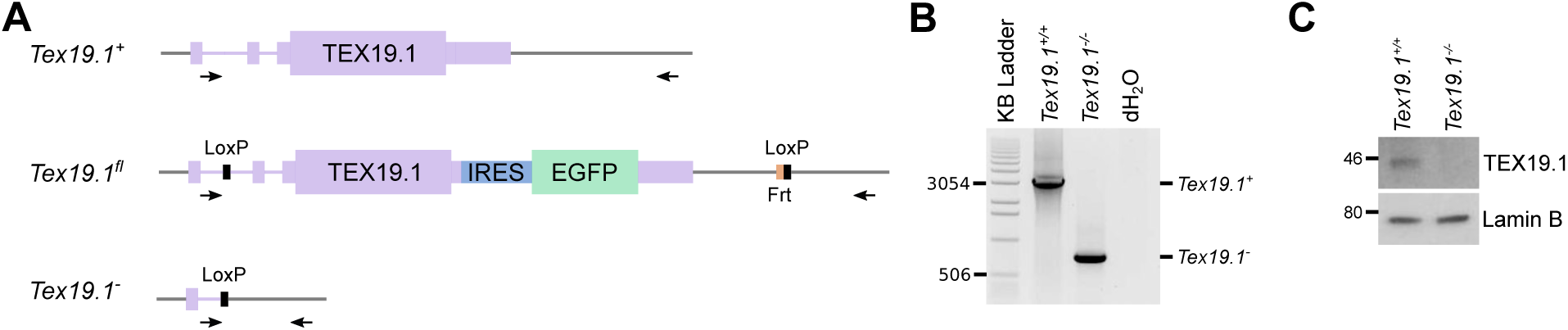
**Generation and Validation Of *Tex19.1***^***−/−***^**ESCs. A.** Schematic diagram showing the *Tex19.1* alleles generated in ESCs. The *Tex19.1* locus is shown in purple, flanking DNA in grey. Introns are shown as lines, exons as rectangles, and the coding region as large rectangles. LoxP sites (black) and an internal ribosome entry site (IRES, blue) coupled to enhanced green fluorescent protein (EGFP, green) were introduced into the locus to generate a *Tex19.1*^*fl*^ allele. This allele also contains an Frt site (orange) left over after excision of a neomycin resistance cassette. After treatment with Cre recombinase the entire *Tex19.1* coding sequence is removed from the *Tex19.1*^*fl*^ allele to generate *Tex19.1*^*-*^. Arrows indicate position of genotyping primers used in panel B. **B.** PCR genotyping of *Tex19.1*^*−/−*^ ESCs. Genomic DNA from *Tex19.*^*+/+*^ ESCs, *Tex19.1*^*−/−*^ESCs or distilled water (dH2O) was used as a template for genotyping PCR using the primers shown in panel A. Migration of selected bands in the KB ladder (Invitrogen) is indicated. **C.**Western blot for TEX19.1 and lamin B in *Tex19.1*^*+/+*^ and *Tex19.1*^*−/−*^ ESCs. MW markers (kD) are indicated.

**Supplementary Figure S8.**
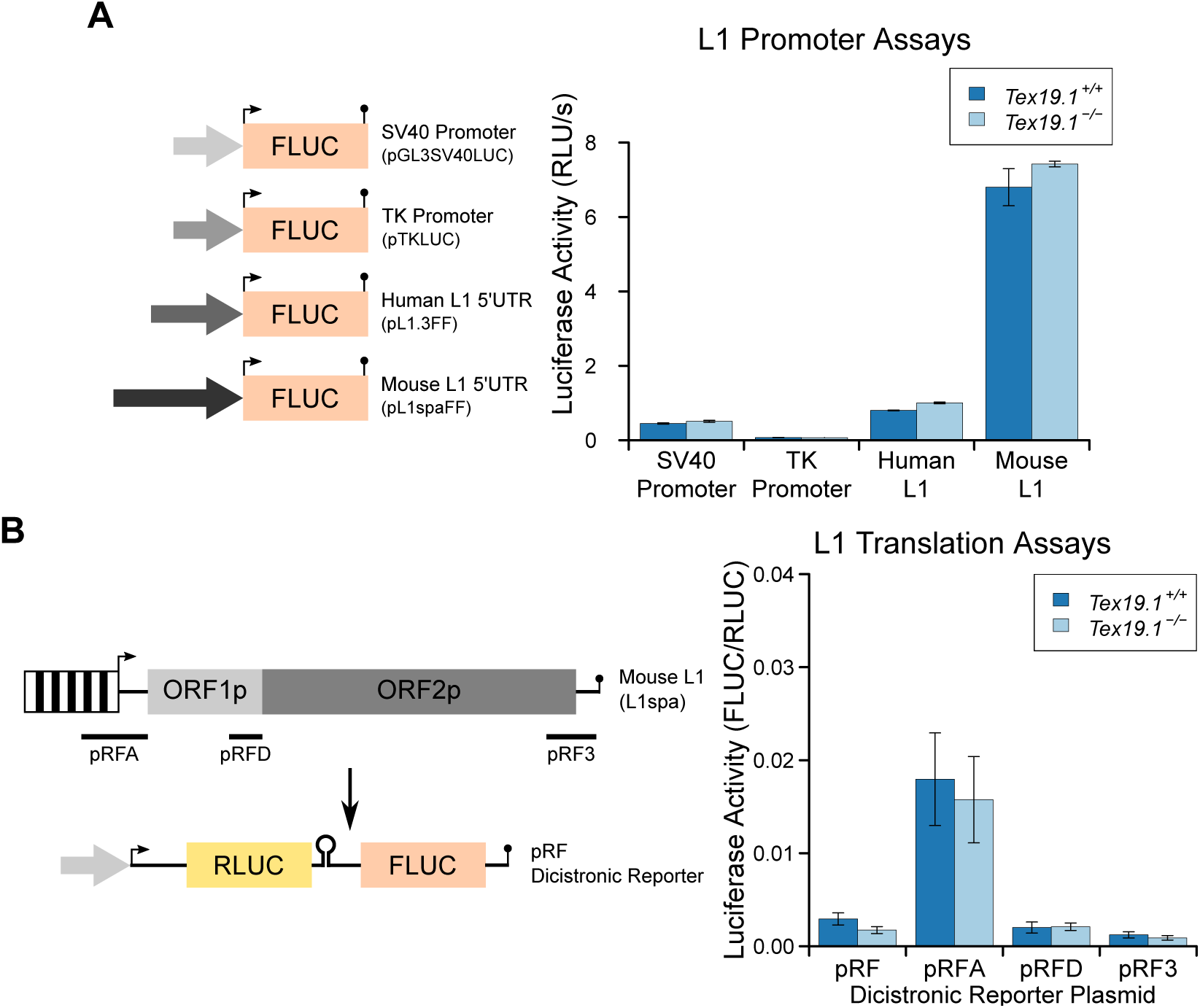
**Loss Of *Tex19.1* Does Not Affect L1 Promoter Or L1 Translation Reporter Activity In Mouse ESCs. A.** Schematic diagram showing promoter-luciferase constructs containing indicated control or L1-derived promoters. Luciferase activity (relative light units per second) of these constructs after transfection into *Tex19.1*^*+/+*^ and *Tex19.1*^*−/−*^ ESCs is shown. Luciferase activity was corrected for transfection efficiencey and normalised to the SV40 promoter construct in control ESCs. Error bars indicate SEM for technical replicates. **B.** Schematic diagram showing translation-luciferase constructs. Regions of L1 (A: 400 bp upstream of ORF1p covering the 5' UTR; D: 200 bp upstream of ORF2p covering the intergenic region; 3: 312 bp from the 3' UTR) inserted in the pRF dicistronic reporter construct (Li et al 2006) were transfected into *Tex19.1*^*+/−*^and *Tex19.1*^*−/−*^ESCs. The pRFD construct contains the ORF2p internal ribosome entry site that binds hnRNPL and nucleolin, cellular factors that restrict L1 (Peddigari et al 2013). Luciferase acivity for these translation-luciferase constructs in *Tex19.1*^*+/−*^ and *Tex19.1*^*−/−*^ ESCs is shown. Firefly luciferase (FLUC) was measured relative to Renilla luciferase (RLUC). Data represents three replicates, error bars represent SEM, there is no statistically significant difference between *Tex19.1*^*−/−*^ESCs and controls in either the promoter or translation assays (*t*-test).

**Supplementary Figure S9.**
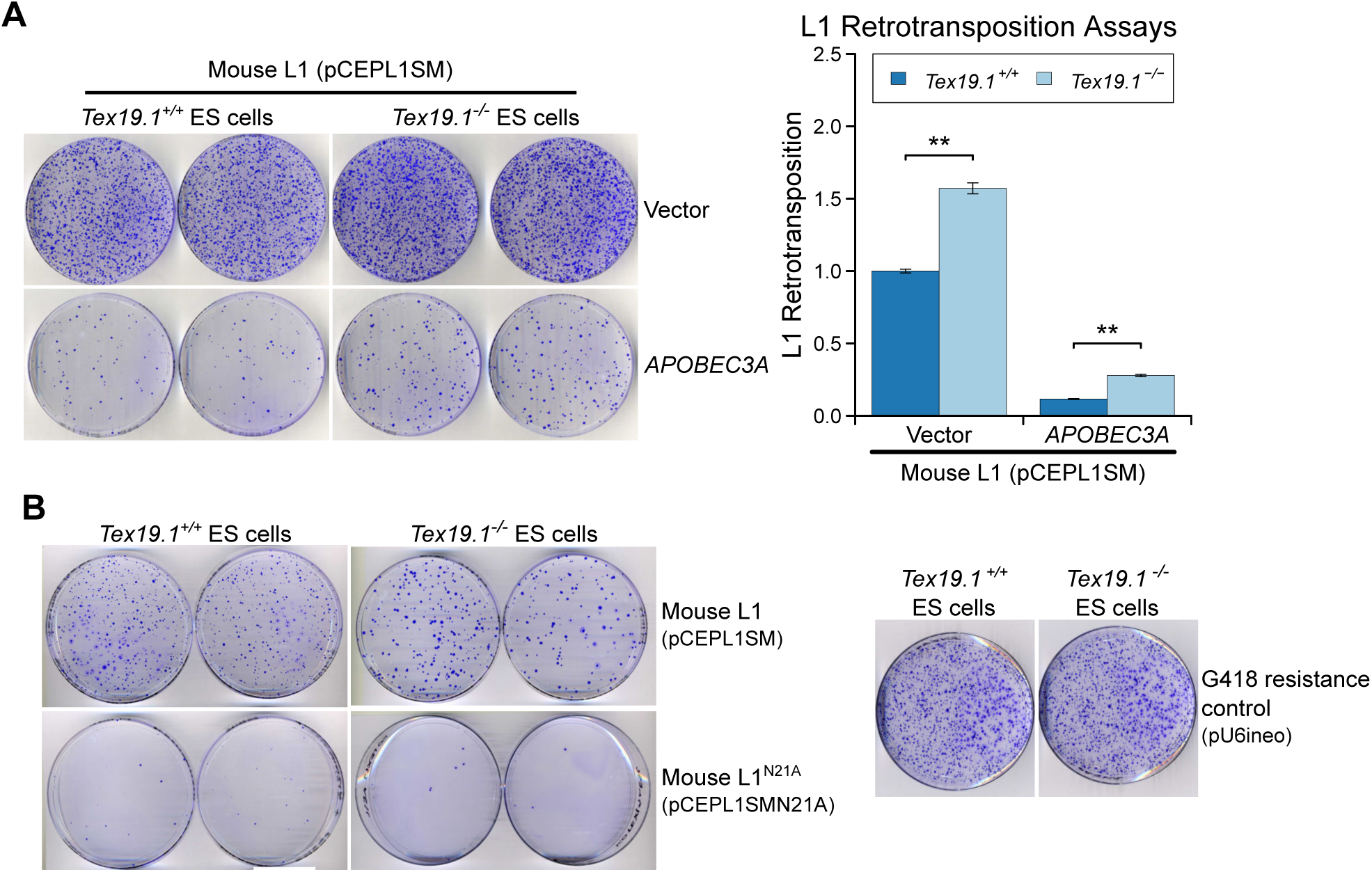
***Tex19.1* Restricts Mobilisation of L1 Reporters in Mouse ESCs. A.** Plates stained with 0.1% crystal violet showing G418-resistant colonies from L1 retrotransposition assays in *Tex19.1*^*+/+*^ and *Tex19.1*^*−/−*^ ESCs. ESCs were co-transfected with a synthetic mouse L1 retrotransposition reporter and either empty vector or the L1 restriction factor *APOBEC3A*. Retrotransposition frequency was calculated relative to *Tex19.1*^*+/+*^ ESCs transfected with empty vector. ** p < 0.01 (*t*-test); error bars indicate SEM. **B.** Additional controls for L1 retrotransposition assays in mouse ESCs. Plates stained with 0.1% crystal violet showing G418-resistant colonies from L1 retrotransposition assays are ahown. Engineered L1 constructs (pCEPL1SMN21A)carrying the N21A mutation in the endonuclease domain of ORF2p that impairs L1 mobilisation (Alisch et al 2006) have greatly reduced retrotransposition in both *Tex19.1*^*+/+*=^ and *Tex19.1*^*−/−*^ ESCs relative to codon-optimisedrelative to codon-optimised L1 (pCEPL1SM). *Tex19.1*^*+/+*^ and *Tex19.1*^*−/−*^ ESCs are able to form colonies when transfected with a control plasmid conferring G418 resistance (pU6ineo).

**Supplementary Figure S10.**
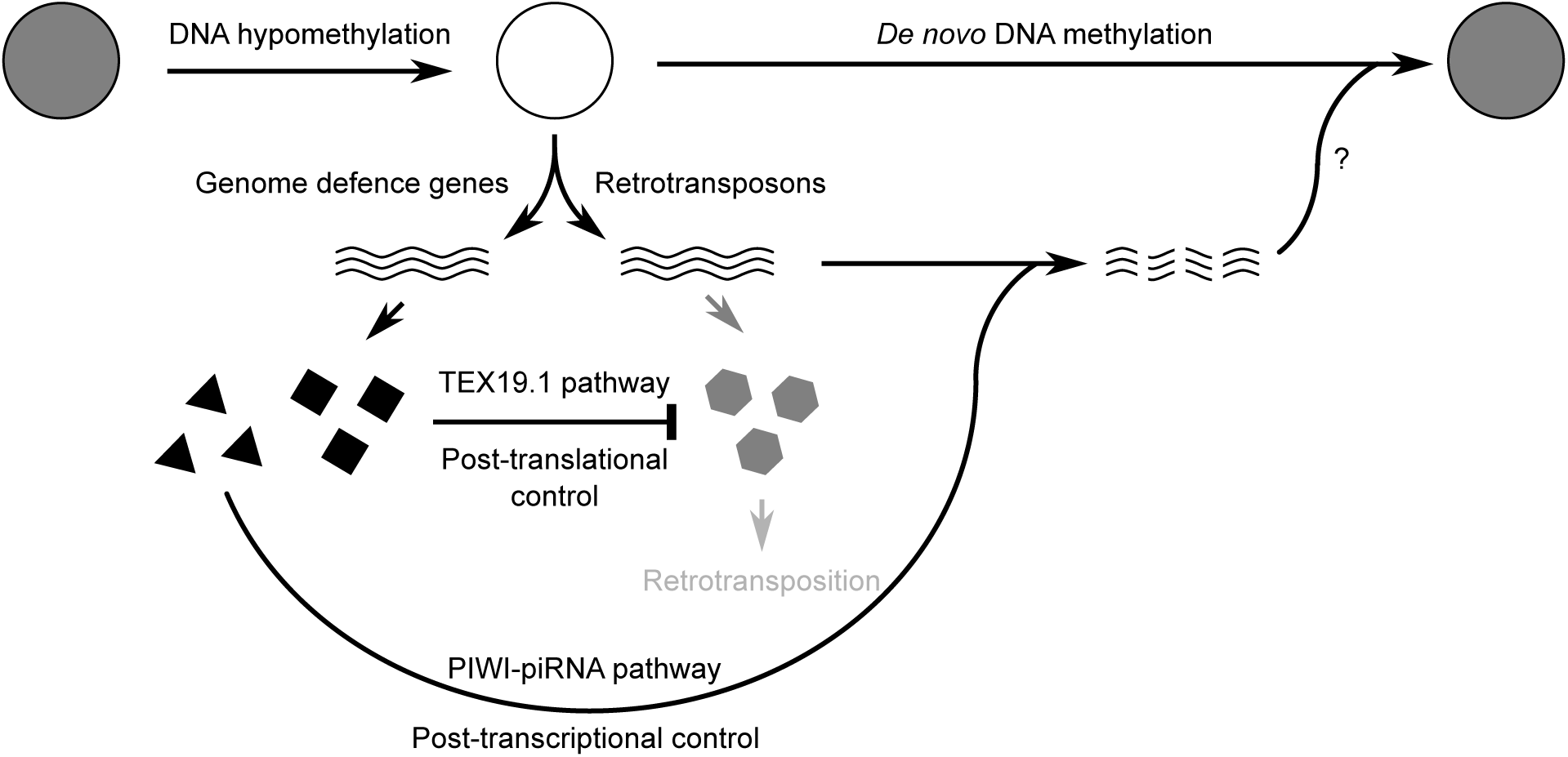
Schematic diagram illustrating how post-translational control mechanisms can contribute to retrotransposon control and genomic stability in hypomethylated male germ cells. RNAs are indicated by wavy lines, proteins by small solid polygons.

